# Patterns of speciation in a parapatric pair of *Saturnia* moths as revealed by Target Capture

**DOI:** 10.1101/2023.07.24.550284

**Authors:** Maria Khan, Mukta Joshi, Marianne Espeland, Peter Huemer, Carlos Lopez Vaamonde, Marko Mutanen

## Abstract

The focus of this study is to understand the evolutionary relationships and taxonomy of widely distributed parapatric species pair of wild silk moths, *Saturnia pavonia* and *Saturnia pavoniella* (Lepidoptera: Saturniidae) in Europe. To address species delimitation challenges associated with many parapatric taxa, target enrichment and mtDNA sequencing was employed alongside phylogenetic, species delimitation, admixture and introgression analyses. The dataset included individuals from both species, two hybrids generated in the lab, as well as individuals from outside the contact zone. Nuclear markers strongly supported both *S. pavonia* and *S. pavoniella* as two distinct species, with the hybrids grouping together as intermediate and separate from both species. However, the maximum likelihood (ML) tree generated from mtDNA sequencing data presented a different picture, showing both taxa to be phylogenetically intermixed. This inconsistency may be attributed to mitonuclear discordance, which can arise from biological factors (e.g., introgressive hybridization or incomplete lineage sorting) or alternatively operational factors (e.g., incorrect species delimitation). We further provide the evidence of past introgression to have taken place, but no evidence of current admixture between the two species. Finally, we discuss our results from evolutionary point of view taking into consideration the past climatic oscillations that has likely shaped the present dynamics between the species. Overall, this study demonstrated the effectiveness of the target enrichment approach in resolving the phylogenetic relationships between closely related parapatric species and providing insights into their taxonomic delimitation.

## Introduction

Species is a fundamental unit of biological diversity. Essentially, all organisms are attributable to a species level, although identification is not always straightforward, and many species remain undescribed and hence unnamed. The fundamental nature of species represents a long-standing debate among the taxonomists, with the semantics of the word ‘species’ being at the core of it. Consequently, over twenty species concepts have been identified in the literature (Hausdorf, 2011; Mallet, 2005; Mayden, 1997). These concepts, despite defining the term in mutually incompatible ways, share one common although broad view - species are separately evolving metapopulation lineages (de Queiroz, 1999). Species delimitation, that is, the process of identifying species-level biological diversity (Carstens et al., 2013) is a crucial aspect of all taxonomic and biodiversity research (Firneno Jr. et al., 2021; Hey, 2001). Despite the importance of species in characterizing biodiversity, delimiting species remains challenging as boundaries between species cannot always be defined unambiguously based on morphological and/or genetic characteristics (Barley et al., 2013). For example, many species tend to exhibit much phenotypic variability across their range, while others appear morphologically indistinguishable despite presence of reproductive isolation between them (Knowlton, 1993). Evolutionary processes such as hybridization, introgression, and incomplete lineage sorting can further complicate the delimitation of species (Harrison & Larson, 2014; Ivanov et al., 2018), especially in case of limited geographic sampling (Chambers & Hillis, 2020). Reaching a consensus over the principles on how species should be delimited under various evolutionary circumstances is a pre-requisite for efficient communication of biodiversity and conservation efforts (Gaston & Spicer, 2004; Tobias et al., 2010).

Climatic oscillations during Pleistocene have shaped the biogeography and distribution of European Lepidoptera to a considerable extent (Schmitt, 2007). These varying geographic modes of distribution can have remarkable effects on species delimitation. As the speciation is often a long and gradual process, the uncertainty of species boundaries is especially pronounced under allopatric and parapatric settings (Joshi et al., 2022; Mutanen et al., 2012; Nosil, 2008). In case of parapatrically distributed taxa, delimitation of species is further complicated by the possibility of admixture and hybridization between the taxa, as the ranges of two taxa may have a narrow zone of overlap with incomplete reproductive barrier (Bull, 1991). It is important to note that observing parapatry in nature is difficult and can be easily regarded as another mode of speciation if debated upon, since parapatric distribution can be the result from either parapatric speciation or secondary contact between populations first diverged in allopatry (Abbott et al., 2013). Parapatric and perhaps incompletely diverged taxa are valuable models to study the speciation process and character displacement through reinforcement (Pfennig, 2016). Some remarkable examples of parapatry displayed in European taxa include butterflies *Melitaea athalia* (Rottemburg, 1775) and *M. celadussa* Fruhstorfer 1910 (Tahami et al., 2021), the Hooded Crow (*Corvus cornix*) and the Carrion Crow (*C. corone*) (Poelstra et al., 2013; Saino et al., 1992; Saino & Villa, 1992), *Pyrgus malvae* Linnaeus 1758 and *P. (malvae) melotis* Duponchel 1832 (de Jong, 1987) as well as the case of *Bombina bombina* and *B. variegata* toads (Yanchukov et al., 2006).

Here, we focused on a parapatric pair of small Emperor moths belonging to Saturniidae family: *Saturnia pavonia* (Linnaeus, 1758) and *Saturnia pavoniella* (Scopoli, 1763). The two species are widely distributed across Europe with *S. pavonia* found in norther and central Europe and *S. pavoniella* in southern and eastern Europe while the situations in Southern France, Iberian and Balkan Peninsulas are still under debate(Huemer & Nässig, 2003; Mazel, 2007). *S. pavoniella* was separated from *S. pavonia* based on the differences in morphological characteristics of the wings and other body parts including the genitalia, as well as the infertility of female F1 hybrids. However, in southern France where *S. pavoniella* populations show a mixture of morphological characters from both taxa (Mazel, 2007). Indeed, both the species have been observed to overlap locally and create a suture zone in southern France, Czech Republic, Italy and Austria. The two taxa are known to show occasional introgression by male hybrids that are fertile (Huemer & Nässig, 2003). Whether this capacity to hybridize has resulted in the disruption of the barrier to gene flow rendering the populations to remain discrete, remains to be studied.

With the increasing availability of genomic-scale data, it is now possible to understand the interplay between different evolutionary processes responsible for formation and distribution of biological diversity (Ferrer Obiol et al., 2023). Although whole genomes are now being sequenced at a relatively fast rate and are potentially ideal source of data for phylogenomic studies, they still remain unavailable for many non-model taxa (Breinholt et al., 2018). Approaches such as target enrichment methods allow for the efficient capture and sequencing of specific regions of the genome, which can be used to infer species boundaries (Mamanova et al., 2010). Further methodological advances have allowed development of species delimitation programs in statistically rigorous frameworks such as Bayesian statistics and multispecies coalescent (Leaché et al., 2014; Yang, 2015; C. Zhang et al., 2018), to infer delimitation from molecular data.

Here, our aim was to shed light on the evolutionary relationships and speciation process of the parapatric and near-cryptic species pair of *S. pavonia* and *S. pavoniella*, and to find out whether they represent a single variable species or two distinct species in terms of biological and phylogenetic species concepts. To address these questions we utilized the target enrichment approach that generates large number of fixed loci across the genome (Joshi et al., 2022; Mayer et al., 2021). In relation to that we further investigated the levels of admixture and traces of historical or ongoing introgression between the two taxa, if any. Motivation to understand this system stems from the desire to find out better justified and more standardized ways to delimit species under complex evolutionary circumstances such as parapatry as this remains largely an unresolved taxonomic problem. The benefits of using a set of standardized genetic markers in establishing a consistent criteria for species delimitation has been discussed in Eberle et al. (2020) and implemented in a set of metazoan taxa in Dietz et al. (2022).

## Materials and Methods

### Taxon sampling and sample preparation

The study dataset involved a total of 27 specimens of genus *Saturnia*, including 11 *S. pavonia*, 13 *S. pavoniella* and 1 *S. josephinae* as an outgroup (Table 1). Additionally, two laboratory hybrid specimens (one male and one female) were also included in the dataset. One of the aims of including lab-reared hybrids was to confirm them as hybrids based on genetic data The specimens were initially identified and assigned to putative species based on morphological characteristics. The samples were collected from nine different countries (Austria, Croatia, Czech Republic, Finland, France, Italy, Slovenia, Spain) from across Europe using the sex pheromone (E6,Z11)-C16Ac synthesized for this study by Till Tolasch (Germany). Lures were prepared at INRAE Orléans from 11-mm red rubber septum lures (Wheaton Scientific, Millville, NJ, USA) loaded with heptane solutions (100 μl, 1 mg/ml) of the synthesized pheromone that had been shipped by courier to INRAE from Tolasch lab in Germany. Loaded septa were placed in a 20-ml glass vial and kept in a freezer when not in use. Lures were sent by post to several colleagues across Europe to attract and collect males for this study (see acknowledgments for list of colleagues and Appendix 1 containing the metadata associated to the vouchers).

**Table 1.** Specimens metadata and dataset statistics.

| Sample ID | Species | Country of Collection | SRA accession | No. of loci recovered | % missing data |
| --- | --- | --- | --- | --- | --- |
| SAT003 | *Saturnia pavonia* | France | SRR23477969 | 453 | 25.40 |
| SAT004 | *Saturnia pavonia* | France | SRR23477968 | 422 | 29.44 |
| SAT006 | *Saturnia pavoniella* | France | SRR23477957 | 409 | 31.58 |
| SAT008 | *Saturnia pavonia* | France | SRR23477949 | 427 | 28.82 |
| SAT009 | *Saturnia pavoniella* | France | SRR23477948 | 456 | 24.32 |
| SAT010 | *Saturnia pavoniella* | France | SRR23477947 | 425 | 29.00 |
| SAT011 | *Saturnia pavoniella* | France | SRR23477946 | 441 | 27.27 |
| SAT012 | *Saturnia pavoniella* | France | SRR23477945 | 456 | 24,00 |
| SAT013 | *Saturnia pavonia* | France | SRR23477944 | 404 | 30.47 |
| SAT020 | *Saturnia pavonia* | Spain | SRR23477943 | 458 | 23.63 |
| SAT021 | *Saturnia pavonia* | Spain | SRR23477967 | 430 | 26.46 |
| SAT022 | Hybrid *pavonia* X *pavoniella* | France | SRR23477966 | 474 | 21.15 |
| SAT023 | Hybrid *pavonia* X *pavoniella* | France | SRR23477965 | 471 | 21.23 |
| SAT024 | *Saturnia pavonia* | Spain | SRR23477964 | 383 | 35.08 |
| SAT026 | *Saturnia pavonia* | Czechia | SRR23477963 | 479 | 20.78 |
| SAT028 | *Saturnia pavoniella* | Czechia | SRR23477962 | 459 | 23.27 |
| SAT031 | *Saturnia pavoniella* | Slovenia | SRR23477961 | 475 | 20.96 |
| TLMF Lep 03073 | *Saturnia pavoniella* | Austria | SRR23477960 | 390 | 33.83 |
| TLMF Lep 30352 | *Saturnia pavonia* | Austria | SRR23477959 | 422 | 28.69 |
| TLMF Lep 30357 | *Saturnia pavoniella* | Austria | SRR23477958 | 407 | 30.73 |
| TLMF Lep 30358 | *Saturnia pavoniella* | Austria | SRR23477956 | 431 | 27,03 |
| TLMF Lep 30361 | *Saturnia pavoniella* | Italy | SRR23477955 | 407 | 30,06 |
| TLMF Lep 30362 | *Saturnia pavoniella* | Italy | SRR23477954 | 425 | 27.95 |
| MM27425 | *Saturnia pavonia* | Finland | SRR23477953 | 447 | 24.91 |
| MM27428 | *Saturnia pavonia* | Finland | SRR23477952 | 433 | 27.93 |
| SAT033 | *Saturnia pavoniella* | Croatia | SRR23477950 | 483 | 18.56 |
| SAT034 | *Saturnia josephinae* | Spain | SRR23477951 | 496 | 20.82 |

The female used to rear the hybrids was a lab reared *S. pavoniella* from the Var region in Southern France and the male was a wild *S. pavonia* from the Loire valley. The crossing was made in a garden in Indre & Loire in central France where the reared *S. pavoniella* female was tied around the base of its two forewings with a piece of 50 – 70 cm of cotton string. The string was itself attached to a shrub to prevent the female from escaping. The female called and mated with one of the wild *S. pavonia* males attracted. The mated *S. pavoniella* female laid eggs and the hybrid caterpillars were reared to obtain F1 hybrid adults.

DNA was extracted from the antenna or thorax of either ethanol preserved or dry specimens. DNA extraction was performed using the QIAGEN DNeasy Blood and Tissue Kit (Hildesheim, Germany) or the Omega Bio-tek E.Z.N.A. Insect DNA Kit (Norcross, United States) following the protocols given by the manufacturers.

### Target Enrichment Library preparation

Target enrichment bait design followed Mayer et al. (2021) where the final probe kit targets 2953 CDS regions in 1753 nuclear genes. This kit was developed with BaitFisher version 1.2.8 (Mayer et al. 2016) and is referred to as the LepZFMK 1.0 kit. Following extraction, the DNA concentration of each sample was measured using the Invitrogen Quant-iT PicoGreen dsDNA (Waltham, United States) quantification kit. Approximately 100 ng of absolute DNA was taken for further processing as per the standard Agilent protocol; for the samples with low DNA yield, approximately 50 ng of absolute DNA was used, depending on the available quantity. For target enrichment protocol the genomic DNA (gDNA) was fragmented using the standard protocols for mechanical DNA shearing on a Diagenode Bioruptor®, to achieve the average fragment size between 150-200 bp. The fragmented DNA was further taken for library preparation using SureSelect XT HS2 library prep kit which included end repair reaction and ligation of adenine residue to the 3’ end of the blunt fragments (A-tailing) to allow ligation of barcoded adaptors. PCR amplification of adaptor-ligated libraries was then performed, after indexing with the Agilent SureSelect XT HS2 primer pairs; we used 8 PCR cycles for all of the samples, irrespective of the amount of absolute DNA taken initially. After the library construction, custom Agilent SureSelect baits (6-11.9 Mb) were used for solution-based target enrichment of a pool containing 8 or 16 libraries. We used LepZFMK 1.0 kit (Mayer et al., 2021), which targets 2953 CDS regions in 1753 nuclear genes, to enrich the gene regions of interest. The hybridisation was performed on each pool with the Agilent SureSelect XT HS2 Target Enrichment Kit ILM module following the manufacturer’s instructions. Enriched libraries were then captured with MyOne Streptavidin T1 beads. These captured libraries were PCR amplified and the final concentration of each captured library was measured using the Invitrogen Quant-iT PicoGreen dsDNA quantification kit. Following the enrichment, pooled libraries were sequenced using the Illumina Nextseq 500 platform (mid output) to generate paired-end 150-bp reads at Biomedicum Functional Genomics Unit (FuGU), Helsinki Institute of Life Science at University of Helsinki.

### Barcoding

For the Mitochondrial DNA (mtDNA) sequencing, the COI gene was amplified using the primers HybLCO and HybHCO. PCR was conducted following the protocols in Wahlberg & Wheat, (2008) (protocol given in supplementary file). The PCR cleanup was then performed using Sephadex columns (Sigma-Aldrich). The purified product was sent to Macrogen Europe for sequencing.

The COI sequences received from Macrogen Europe were aligned using ClustalW as implemented in MEGA X software v10.0.5 (Kumar et al., 2018) and BioEdit 7.2.5 sequence alignment editor (Hall, 1999). The reference sequence against which the other sequences were aligned was retrieved from BOLD Systems 4.0 database (http://www.boldsystems.org). This reference specimen was one of those included in the sample specimens. For the specimens which failed the sequencing, we extracted the barcode region from the target enrichment data using *exonerate* ver. 2.4.0 (Slater & Birney, 2005), with the alignment of individuals that were successfully barcoded as a query and our de-novo assembled target enrichment assemblies as a target (Details of the command are given in supplementary file). These extracted barcodes were aligned with the rest of the barcodes using Muscle aligner on EMBL-EBI server (https://www.ebi.ac.uk/Tools/msa/muscle/) with default parameters. The alignment was manually checked in Geneious Prime (https://www.geneious.com/prime/) before proceeding for phylogenetic analysis.

To visualize divergence patterns in COI, phylogenetic trees of the aligned COI sequences were generated using IQTREE ver. 2.0.3 (Minh et al., 2020), using the best-fit model estimated by the program using ModelFinder (Kalyaanamoorthy et al., 2017) and ultrafast bootstrapping with 1000 replicates. We used -bnni option to reduce the impact of model violation and overestimated branch support. The phylogenetic analysis was run 20 times for each of the study species to assess the congruence among the tree searches. The trees with the highest likelihood were visualized in FigTree v1.4.4 (available from tree.bio.ed.ac.uk/software/figtree/).

### Target Enrichment bioinformatics

The demultiplexed data after being checked for quality was processed using the TEnriAn pipeline (Mayer et al., 2021). During the first step of TEnriAn workflow, adaptor sequences and low-quality bases were removed from the demultiplexed data using fastp (Chen et al., 2018). The reads were then assembled using Trinity version 2.9.0 (Grabherr et al., 2011). We skipped the Contamcheck step during the assembly, where the data is checked and filtered for potential cross-contaminations, using ‘all against all’ blast searches as it was observed to be removing lot of useful data due our samples being very closely related. Instead, we used the original assembled files before contamination checking and used them as an input for the Orthograph step. During this step, clusters of orthologous loci are inferred, using a pre-generated reference database. In our case, this reference database was generated using the coding gene alignments of 5 species – *Bombyx mori, Danaus plexippus, Heliconius melpomene, Melitaea cinxia, Papilio glaucus.* Next, the orthologous clusters were aligned, using reference Hidden Markov Models (HMMs) in hmmalign (part of the HMMER package, http://hmmer.org), and the alignments were filtered various criteria such as coverage, outliers, following Mayer et al. (2021). We further removed the loci having GC content of higher than 60%, as those can have a negative effect on downstream phylogenetic analysis.

### SNP Calling

To perform population genetic analyses, we also did variant calling on the target enrichment dataset. We used the ingroup sample with the highest number of orthologous clusters inferred overall (during orthology assessment step in the TEnriAn workflow) as a reference for SNP calling. The raw cleaned data (i.e., after removing adaptors and low-quality bases) was mapped against this reference using with the BWA-MEM algorithm in bwa 0.7.17 (available from bio-bwa.sourceforge.net) with the minimum seed length set to 30.

SNPs were then extracted from the sorted BAM files generated during mapping by using the samtools *mpileup* option piped together with the bcftools call option. Then filtering was done using bcftools to remove indels and multiallelic SNPs and keeping only biallelic SNPs. The number of SNPs obtained after this filtering step were 10,265.

### Phylogenetic Analyses

The filtered alignments from the TEnriAn workflow were concatenated using FASConcat-G available at https://github.com/PatrickKueck/FASconCAT-G. This concatenated data was used to infer Maximum Likelihood tree using IQTREE 2.0.3 (Minh et al., 2020). In order to partition the data according to loci, we set up partitioned analysis using IQTREE with option -m TESTMERGEONLY to resemble PartitionFinder (Chernomor et al., 2016) and the rcluster algorithm (Lanfear et al., 2014) with the rcluster percentage set to 10, under the AICc criterion. The best partitioning scheme was then used as an input to set up a partitioned analysis in IQ-TREE. We used the ultrafast bootstrap approximation with 1000 replicates (Hoang et al., 2018). To further reduce the impact of severe model violations, the -bnni option was used along with UFBoot. We also performed a SH-like approximate likelihood ratio test (Guindon et al., 2010) with 1000 replicates using the -alrt option. Each analysis was run 20 times to assess the congruence among the tree searches. The tree with the highest likelihood was visualised in Figtree.

Generally, in phylogenomic datasets with multiple loci, some amount of discordance is expected among the gene trees. For this reason, it is useful to build so called ‘true species tree’, reflecting the correct relationships among the species of interest. For this, we used ASTRAL-III v. 5.7.4 (C. Zhang et al., 2018), Since it is statistically consistent under multispecies coalescent model and can handle incomplete lineage sorting in the dataset. A set of gene trees were inferred using IQTREE version 2.0.3 (Minh et al., 2020) where IQTREE was prompted to perform model selection (using ModelFinder) and tree inference separately for each locus. To improve the ASTRAL results further, the gene trees were filtered using program TreeShrink (Mai & Mirarab, 2018) before running ASTRAL to detect and prune any abnormally long branches. The program was run in default ‘per-species’ mode with a false positive tolerance rate (α) set to 0.05. The resulting shrunk output gene trees were used as an input for ASTRAL, which generated a species tree along with the quartet score. The tree was visualized in FigTree v1.4.4.

### Population Genetics

To get a notion of the possible number of genetic clusters and related species in the dataset we performed Principal Component Analysis (PCA) on the target enrichment SNP dataset. The PCA was ran using *dudi.pca* function in the ‘adegenet package’ (Jombart & Ahmed, 2011) in R studio v1.4.1106. We also calculated the pairwise F_ST_ from the same SNP dataset using R package *hierfstat* (Goudet, 2005).

STRUCTURE analysis (Pritchard et al., 2000) was also conducted on the target enrichment SNP dataset to infer population structure as well as admixture patterns. We used unlinked SNPs (as one of the assumptions of STRUCTURE includes that the loci are at HW equilibrium), and for this SNP data was filtered using a custom Perl script to keep only one SNP per scaffold. The final number of SNPs after this filtering was 1,831. The analysis was done assuming five clusters, that is for K=1 to K=5, each with 10 replicate runs. We used Clustering Markov Packager Across K or CLUMPAK ver. 1.1 (Kopelman et al., 2015) to align the cluster assignments across all 10 replicates for K = 2, K = 3, and K = 4 and Distruct program to show barplots for the K populations. Structure Harvester (Earl & vonHoldt, 2012) was used to obtain optimal number of genetic clusters (*K*) using ΔK method with 500,000 generations for the Markov chain and a value of 100,000 as burn-in. The program produces a graph based on the Evanno method (Evanno et al., 2005), between K cluster and DeltaK along X and Y axis respectively, which shows the K with the highest DeltaK along the axes.

### Species Delimitation Analyses

To perform species delimitation analyses, we conducted Trinomial distribution analysis tr2 (Fujisawa et al., 2016), in which the distribution of rooted triplets is modeled to find the congruence between gene trees under multispecies coalescent framework. In short, it measures the incongruence of tree topologies within species and congruence between species. We tested a two-species hypothesis compared to the null model (assuming a single species) and obtained the results based on calculation of –log (posterior probability).

Additionally, we calculated the Genealogical Divergence Index (gdi) (Jackson et al., 2017) for the genetically distinct clades identified from phylogenetic analyses, with BPP using the approach described in Leaché et al. (2019). For this, we used our complete target enrichment dataset including all the loci. We first conducted a full BPP analysis (A11), where species tree and species delimitation are jointly estimated. We observed that BPP consistently inferred delimitation where all the populations were assigned to distinct species with posterior probability of 1, irrespective of prior combinations used. This is likely caused by the likelihood problem and should be explored in detail with empirical data. Next, we conducted BPP analysis where species delimitation and species tree are fixed, to infer theta and tau values for each node corresponding to the distinct clades in the tree (BPP A00 analysis). We used beta prior values 0.004 for theta (population size) and 0.002 for tau (divergence times) respectively and tree inferred by ASTRAL as a guide tree. The theta values were estimated using Bayesian analysis, by specifying ‘e’ option in A00 analysis, which was conducted 2 independent times using MCMC chain length of 1000000 and burn-in of 100000. The convergence of the two runs was assessed in Tracer v1.7.2 (Rambaut et al., 2018) and the converged runs were concatenated to generate posterior distributions for the multispecies coalescent parameters. The gdi was calculated as

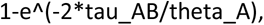

where tau_AB is the inferred divergence time of the clade from its sister group, and theta_A is the inferred population size for the clade. We determined the final theta and tau values as the median theta and tau values from the converged run. The gdi value of less than 0.2 supports the conspecificity of sister clades, values between 0.2 and 0.7 indicates the so-called ‘grey zone’, while gdi value of > 0.7 supports them being a distinct species (Jackson et al., 2017).

### Isolation by distance

We tested for isolation by distance (IBD) using a Mantel test between a matrix of genetic distances and matrix of geographic distances using the *mantel.randtest* function in R package *adegenet*. This test finds the correlation between individual Edwards’ genetic distances and Euclidean geographic distances (Mantel, 1967). Based on the simulated p-value, we determined whether isolation by distance was significant (p < 0.05).

### D-statistics

We used the program Dsuite (Malinsky et al., 2021) to calculate the Patterson’s D-statistics to test for the presence of historical introgression in our dataset. This test was first introduced in Green et al. (2010) to detect the presence of Neandertal ancestry in modern humans, with further theoretical improvements and applications in Reich et al., (2010) and Durand et al., (2011). This test computes the excess of alleles in one species (usually denoted as P3) in relation to two sister species (P1 and P2), and a fourth outgroup species as a reference. For this, we used a vcf file containing multiallelic SNPs as an input. The Dsuite program uses only biallelic SNPs by default, therefore eliminating the need to filter multiallelic SNPs file beforehand. Since we did not have three populations (excluding outgroup), we ran this program assuming every specimen as a distinct species/population, excluding hybrids from the analysis. D-statistics was calculated using *Dtrios* command, which calculates ABBA-BABA proportions and f4-ratio statistics for all possible trios of populations/species. We also performed *f*-branch (*f*b) test which calculates *f*-branch statistic values for each branch on the input tree, including internal branches (Malinsky et al., 2021) and is useful in guiding the interpretation of correlated f4-ratio results. We used the tree inferred from ASTRAL as an input hypothesis tree, to calculate the *f*-branch statistic.

## Results

### Overview of the Dataset

For our genomic dataset the average number of informative loci retained was 439 (SD=29.7, Table 1), with an average amount of missing data of 26.42% (SD=0.043, Table 1). For our Mitochondrial DNA (mtDNA) dataset, 23 specimens were successfully sequenced, and for the rest we used the barcodes extracted from target enrichment dataset for further analysis.

### Phylogenetic Analyses

The preliminary specimen identification was done based on the morphological characteristics. The ML tree obtained from mitochondrial COI dataset (Fig. 2), did not show reciprocal monophyly, and the taxa seemed to be phylogenetically intermixed. However, five specimens of *S. pavoniella* formed a sister group to the rest specimens, suggesting that at least this species is having species-specific haplotypes.

**Figure 1:**
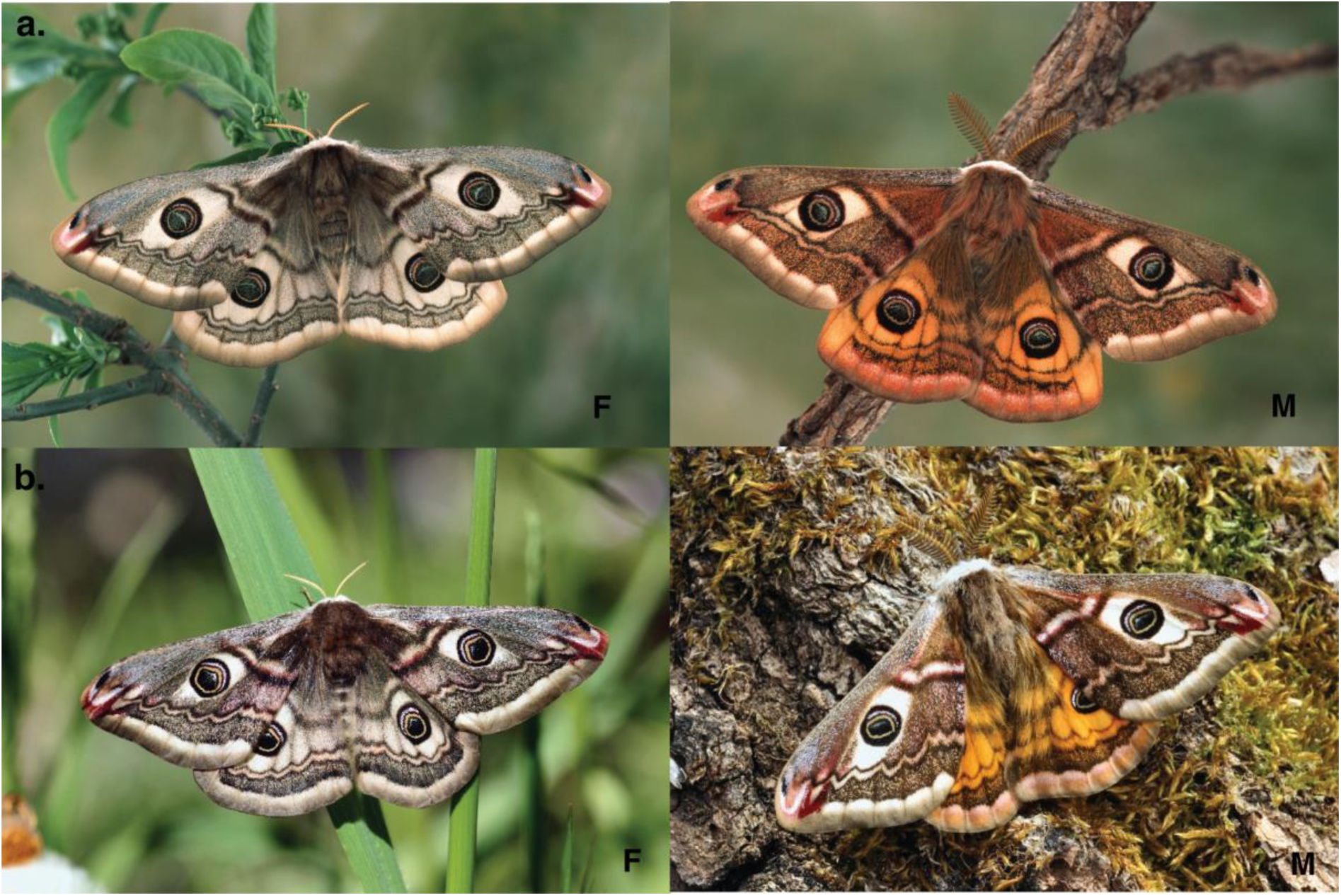
Representative images of live wild male and female (a) *S. pavoniella* and (b) *S. pavonia* specimens

**Figure 2:**
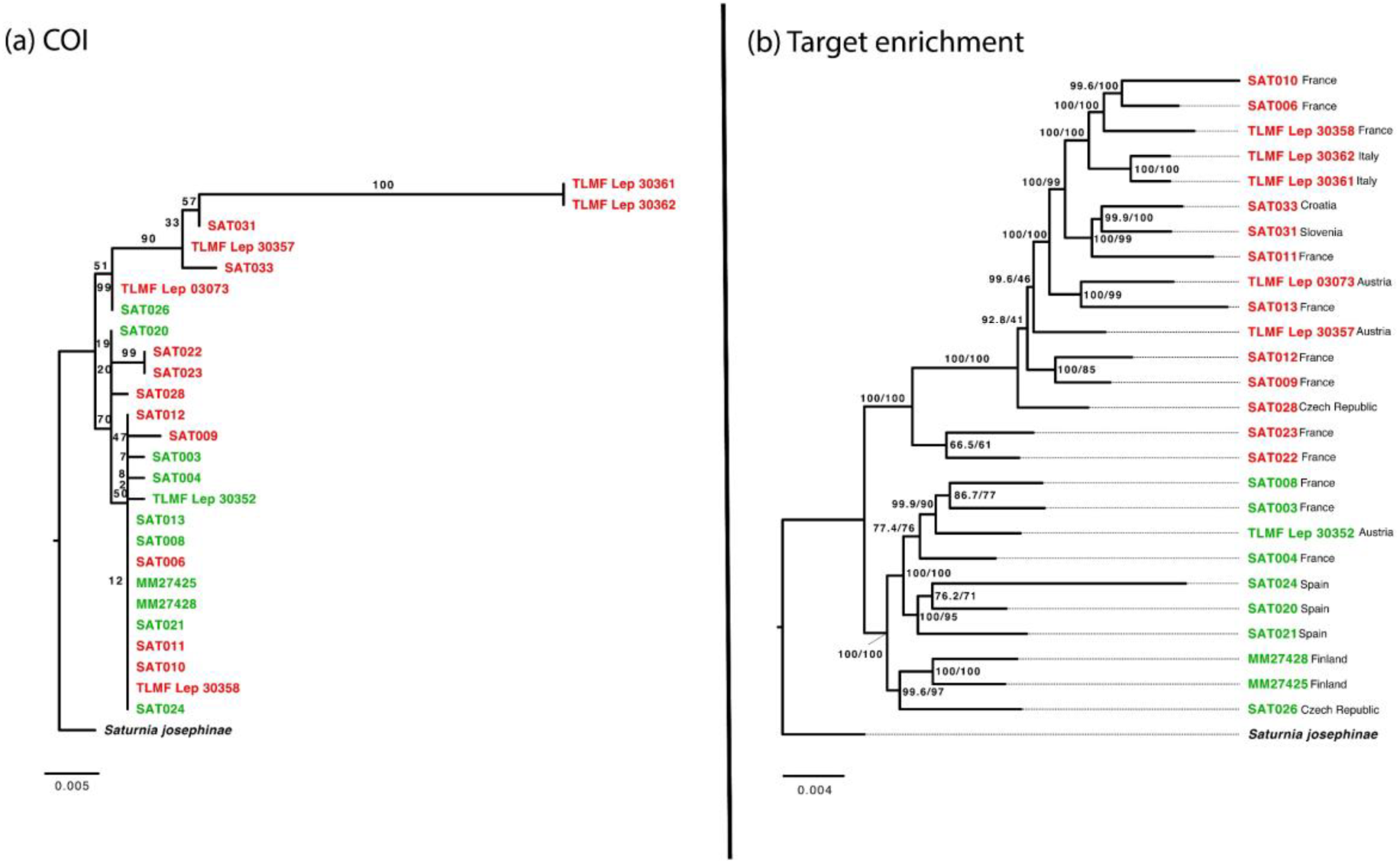
Maximum likelihood tree inferred using barcode data (on left) and target enrichment data (on right). *S. pavonia* individuals are indicated by green color and *S. pavoniella* by red color.

The ML tree from the target enrichment dataset showed two separate and strongly supported clusters corresponding to *S. pavonia* (indicated by green, Fig 2) and *S. pavoniella* (indicated by red, Fig 2), with SH-aLRT/UFBoot support values 100/100 and 100/99 respectively. The hybrids did not show monophyly with either one of two clusters, and were observed to be paraphyletic with respect to *S. pavoniella* clade. One individual from *S. pavonia* (ID SAT024) has an unstable longer branch length which could be potentially caused by errors during alignment step, or alternatively due to sequencing errors that were not filtered out at the bioinformatics step.

We observed that trees obtained from input dataset filtered using TreeShrink did not differ from the one obtained using data without filtering. Both the trees (filtered and unfiltered) correspond to the ML tree generated from our genomic dataset (Fig. 3 and Fig. S5). Similar to the ML tree, *S. pavonia* and *S. pavoniella* were separated into two clusters with quartet support values of 0.99 and 1, respectively. However, unlike the ML tree, the hybrids were grouped together in ASTRAL tree (with quartet support value of 0.83). We further observed that in both COI and ASTRAL tree, two individuals from Italy, TLMF Lep 30361 and TLMF Lep 30362 were placed on an internal branch longer than the rest of the individuals.

**Figure 3:**
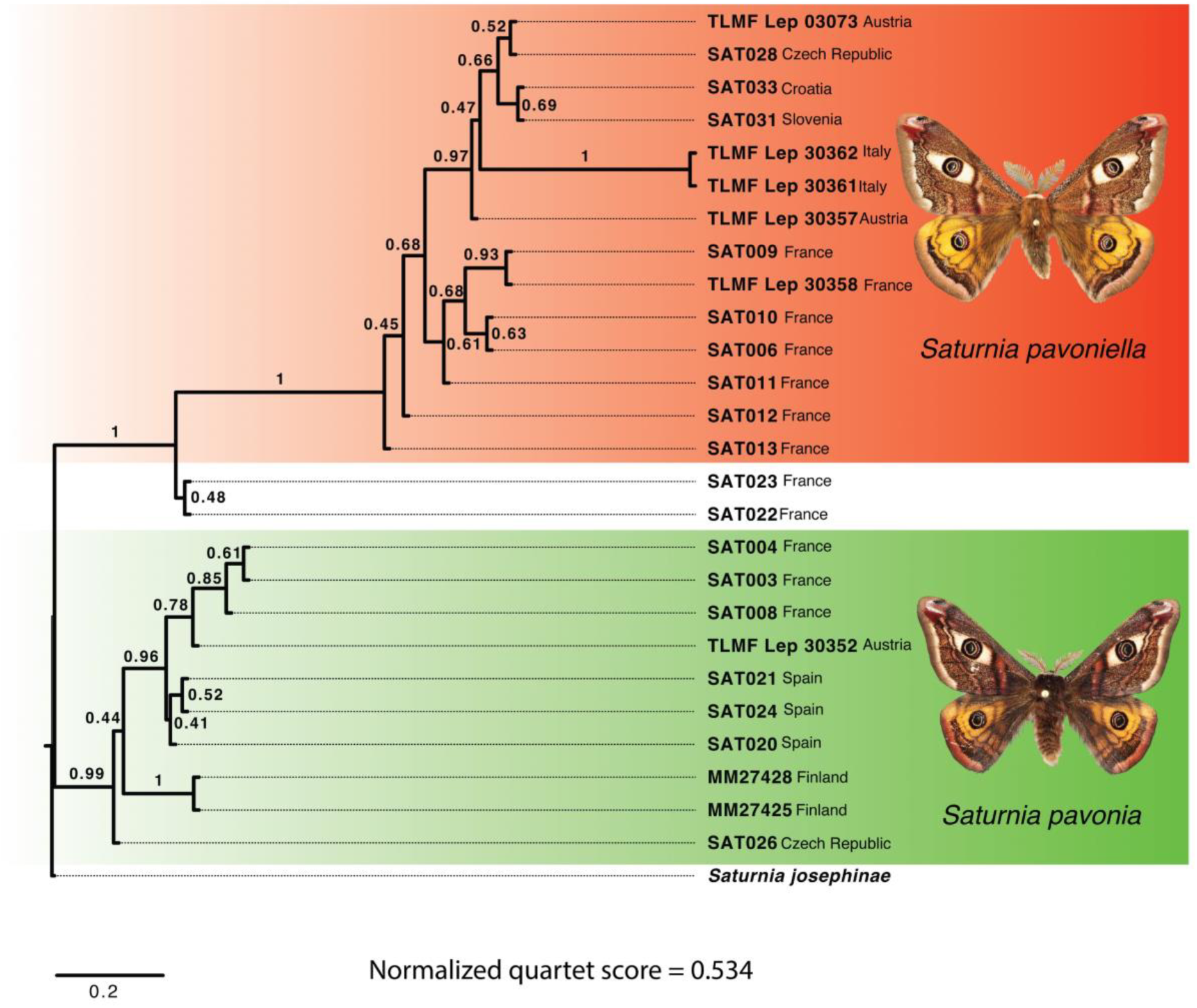
ASTRAL tree inferred for *Saturnia.* Numbers on the branches indicate quartet support values. Both specimens SAT022 (male) & SAT023 (female) are lab reared F1 hybrids obtained from the following crossing: pavoniella female * pavonia male

### PCA and F_ST_

PCA based on the SNP dataset resulted in two genetically distinct clusters for *S. pavonia* and *S. pavoniella* (Fig 4). The two individuals that are situated between these two clusters in the figure are hybrids, that appear intermediate and stay separated from both clusters (Fig. 4). Three *S. pavonia* specimens with sample IDs. SAT026, MM27425 and MM27428 are separated from the rest of the *S. pavonia* individuals mainly along PC2, indicating the presence of genetic differentiation to some extent in these 3 individuals. The pairwise F_ST_ between the two species was calculated as 0.345.

**Figure 4:**
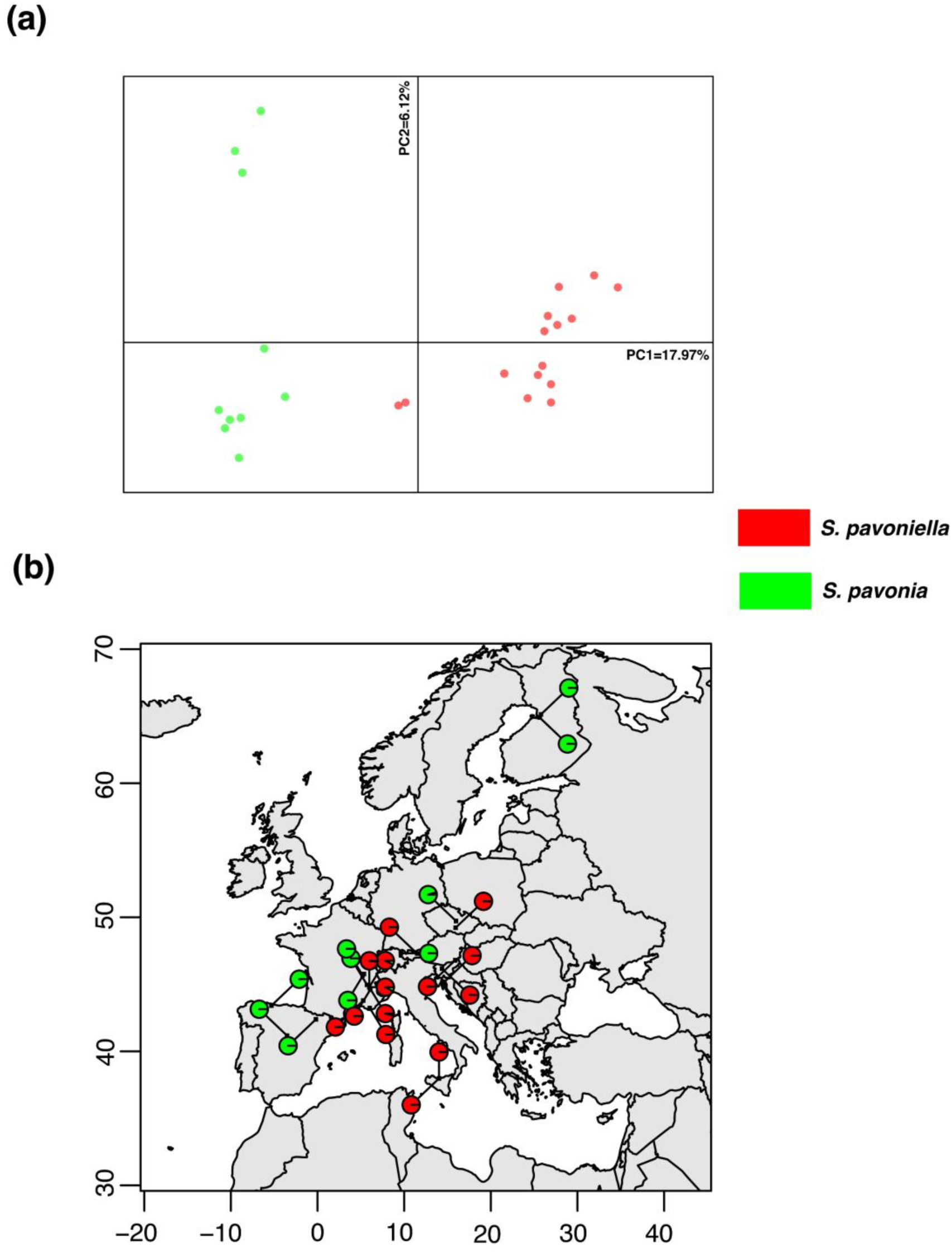
a) PCA for *Saturnia* SNP dataset b) Pie charts based on membership coefficient matrix at K=2 mapped on geographic coordinates

### Genomic admixture and isolation by distance

The analysis of genomic admixture using STRUCTURE produced the highest likelihood estimate for K=2 clusters for the target enrichment SNP dataset as can be seen in Table S1. The CLUMPAK analysis of STRUCTURE results gave bar plots that show cluster assignment from K=2 to K=4 (Supplementary figure S1). The results do not suggest admixture between the two parapatric taxa. The only admixture that was observed was in the hybrids that were included in the study. From the geographic distribution of genomic admixture, it can be observed that both species have maintained genetic separation, despite range overlap in the hybrid zone (Fig. 4). Only a slight admixture was observed (from K=3) in a few *S. pavonia* and *S. pavoniella* individuals (supplementary figure S4).

The isolation by distance was observed to be non-significant (p-value > 0.05, supplementary figure S3)

### Species delimitation

The tr2 species delimitation analysis gave log-likelihood scores based on posterior probability for the null model (assuming *S. pavonia* and *S. pavoniella* as a single species), and model1 (assuming *S. pavonia* and *S. pavoniella* as two different species). The calculated values were - 231271.46 and -7339.42 for the null model and model1 respectively, showing that the likelihood is higher for model1 which is the two-species model. The genealogical divergence index (gdi) value for divergence of *S. pavonia* clade from *S. pavoniella* was calculated as 0.277, suggesting the distinct species status of the two clades being ambiguous.

### Introgression

Out of 2024 trios analyzed, high D-statistics values with significant P-values were observed for about 515 trios (D statistics > 0.25, supplementary table S2). From these, we further separated the trios where both P1 and P2 are represented by same species, after which we obtained about 182 trios where both P1 and P2 are represented by *S. pavonia* and P3 by *S. pavoniella* and about 223 trios where P1 and P2 are represented by *S. pavoniella* and P3 by *S. pavonia*. Out of these, significantly high D-statistic value (>=0.4) was observed for about 53 trios (Table 2) and additionally an elevated f4 ratio (supplementary figure S2). However, Malinsky et al. (2021) showed that the D and *f*_4_ -ratio statistics are correlated and that a significantly elevated result for a trio does not necessarily indicate that the populations are involved in gene flow event. But, we observed stronger *f*-branch signal, i.e., between 5-10% for several branches including pavonia7 (SAT026) and an entire *pavoniella* clade as well as pavoniella14 (SAT013) and several *pavonia* branches (Fig. S2c), indicating past introgression incidents between the two species.

**Table 2:** D-statistics summary for SNP dataset.

| P1 | P2 | P3 | Dstatistic | Z-score | p-value | f4-ratio | BBA A | ABB A | BAB A |
| --- | --- | --- | --- | --- | --- | --- | --- | --- | --- |
| SAT020 | SAT026 | TLMF Lep 30362 | 0,49 | 10,19 | 2,30E-16 | 0,14 | 213,25 | 195,75 | 67 |
| TLMF Lep 30361 | SAT021 | SAT020 | 0,49 | 8,69 | 2,30E-16 | 0,14 | 613,25 | 69,25 | 23,75 |
| SAT020 | SAT026 | SAT031 | 0,49 | 9,39 | 2,30E-16 | 0,16 | 216,75 | 209,25 | 72,5 |
| SAT020 | SAT026 | TLMF Lep 30357 | 0,49 | 9,19 | 2,30E-16 | 0,15 | 200,25 | 186 | 64,5 |
| TLMF Lep 30361 | SAT012 | SAT021 | 0,48 | 6,36 | 1,97E-10 | 0,15 | 646 | 63,25 | 22 |
| TLMF Lep 30362 | SAT012 | SAT021 | 0,48 | 6,41 | 1,42E-10 | 0,15 | 658,75 | 66,25 | 23,25 |
| SAT020 | SAT026 | TLMF Lep 30361 | 0,48 | 9,16 | 2,30E-16 | 0,14 | 212,5 | 189,75 | 67,25 |
| SAT020 | SAT026 | TLMF Lep 03073 | 0,46 | 8,60 | 2,30E-16 | 0,15 | 195,25 | 185,75 | 68,25 |
| SAT020 | SAT026 | SAT033 | 0,46 | 9,24 | 2,30E-16 | 0,14 | 220,75 | 202,25 | 74,5 |
| TLMF Lep 30361 | SAT013 | SAT020 | 0,46 | 7,47 | 7,75E-14 | 0,13 | 566,75 | 66,5 | 24,75 |
| TLMF Lep 30362 | SAT013 | SAT020 | 0,45 | 7,11 | 1,13E-12 | 0,13 | 573,5 | 65,75 | 25 |
| TLMF Lep 30362 | SAT013 | SAT021 | 0,44 | 5,53 | 3,29E-08 | 0,13 | 615,375 | 58,625 | 22,625 |
| TLMF Lep 30362 | SAT012 | SAT020 | 0,44 | 6,83 | 8,51E-12 | 0,13 | 625 | 71 | 27,5 |
| TLMF Lep 30361 | SAT013 | SAT021 | 0,44 | 5,25 | 1,53E-07 | 0,13 | 605,625 | 58,875 | 22,875 |
| TLMF Lep 30361 | SAT012 | SAT008 | 0,44 | 5,25 | 1,51E-07 | 0,13 | 591,875 | 73,875 | 28,875 |
| TLMF Lep 30361 | SAT013 | TLMF Lep 30352 | 0,43 | 5,78 | 7,40E-09 | 0,12 | 528,125 | 69,875 | 27,625 |
| SAT021 | MM27425 | TLMF Lep 03073 | 0,43 | 8,46 | 2,30E-16 | 0,12 | 176,875 | 163,125 | 64,625 |
| SAT021 | MM27425 | TLMF Lep 30362 | 0,43 | 8,80 | 2,30E-16 | 0,11 | 189 | 163,25 | 65 |
| TLMF Lep 30361 | SAT013 | SAT008 | 0,43 | 6,54 | 6,27E-11 | 0,13 | 547,375 | 70,625 | 28,375 |
| SAT021 | MM27428 | TLMF Lep 03073 | 0,43 | 5,74 | 9,71E-09 | 0,11 | 179,125 | 156,375 | 62,875 |
| TLMF Lep 30362 | SAT012 | SAT008 | 0,43 | 5,08 | 3,82E-07 | 0,14 | 604,875 | 81,125 | 32,625 |
| TLMF Lep 30362 | SAT013 | TLMF Lep 30352 | 0,43 | 5,87 | 4,48E-09 | 0,12 | 536,875 | 68,625 | 27,625 |
| SAT008 | SAT026 | SAT028 | 0,42 | 7,10 | 1,21E-12 | 0,15 | 218,875 | 197,375 | 80,125 |
| TLMF Lep 30362 | SAT028 | SAT020 | 0,42 | 3,12 | 0,00182564 | 0,08 | 685,25 | 43,5 | 17,75 |
| SAT020 | SAT026 | SAT009 | 0,42 | 9,40 | 2,30E-16 | 0,13 | 216,875 | 192,875 | 78,875 |
| TLMF Lep 30361 | SAT013 | SAT003 | 0,42 | 5,31 | 1,10E-07 | 0,13 | 532,875 | 75,375 | 30,875 |
| SAT033 | SAT012 | SAT021 | 0,42 | 4,81 | 1,48E-06 | 0,13 | 694,5 | 62,75 | 25,75 |
| TLMF Lep 30362 | TLMF Lep 30358 | SAT021 | 0,42 | 3,41 | 0,00065793 | 0,10 | 703,25 | 47,5 | 19,5 |
| SAT033 | SAT012 | SAT020 | 0,42 | 6,35 | 2,09E-10 | 0,13 | 663 | 71,25 | 29,25 |
| TLMF Lep 03073 | SAT013 | SAT021 | 0,42 | 4,59 | 4,36E-06 | 0,13 | 603,125 | 58,625 | 24,125 |
| SAT021 | MM27425 | SAT028 | 0,42 | 10,70 | 2,30E-16 | 0,12 | 179,625 | 176,125 | 72,625 |
| TLMF Lep 03073 | SAT013 | SAT003 | 0,42 | 5,25 | 1,48E-07 | 0,13 | 524,125 | 74,875 | 30,875 |
| SAT033 | SAT012 | SAT008 | 0,41 | 6,08 | 1,19E-09 | 0,13 | 640,625 | 80,625 | 33,375 |
| SAT021 | MM27428 | TLMF Lep 30362 | 0,41 | 5,25 | 1,48E-07 | 0,10 | 195,25 | 153,75 | 64 |
| TLMF Lep 30361 | TLMF Lep 30358 | SAT020 | 0,41 | 4,99 | 5,97E-07 | 0,08 | 659,875 | 46,375 | 19,375 |
| SAT020 | SAT026 | TLMF Lep 30358 | 0,41 | 9,68 | 2,30E-16 | 0,13 | 195,125 | 186,625 | 78,125 |
| TLMF Lep 30361 | SAT012 | SAT004 | 0,41 | 5,21 | 1,91E-07 | 0,12 | 566,875 | 70,125 | 29,375 |
| SAT021 | MM27425 | TLMF Lep 30361 | 0,41 | 7,08 | 1,45E-12 | 0,10 | 185,875 | 159,125 | 66,875 |
| TLMF Lep 30361 | SAT011 | SAT004 | 0,41 | 4,99 | 5,97E-07 | 0,12 | 530,125 | 67,375 | 28,375 |
| SAT008 | SAT026 | SAT031 | 0,41 | 7,05 | 1,76E-12 | 0,13 | 237,125 | 194,375 | 81,875 |
| SAT021 | MM27428 | SAT028 | 0,41 | 5,84 | 5,07E-09 | 0,12 | 186,125 | 164,375 | 69,375 |
| SAT021 | SAT004 | TLMF Lep 03073 | 0,41 | 5,94 | 2,79E-09 | 0,10 | 204,375 | 137,625 | 58,125 |
| TLMF Lep 30361 | SAT013 | SAT004 | 0,41 | 5,54 | 3,04E-08 | 0,12 | 528,625 | 65,375 | 27,625 |
| SAT021 | SAT004 | SAT028 | 0,40 | 7,21 | 5,77E-13 | 0,10 | 200,125 | 147,375 | 62,625 |
| SAT004 | SAT026 | SAT031 | 0,40 | 6,40 | 1,58E-10 | 0,12 | 226,125 | 174,875 | 74,375 |
| TLMF Lep 30362 | SAT013 | SAT008 | 0,40 | 6,13 | 8,66E-10 | 0,12 | 554,125 | 66,625 | 28,375 |
| SAT021 | MM27428 | TLMF Lep 30357 | 0,40 | 5,36 | 8,43E-08 | 0,11 | 175,5 | 158,75 | 67,75 |
| TLMF Lep 30357 | SAT012 | SAT021 | 0,40 | 5,34 | 9,42E-08 | 0,13 | 644,125 | 61,125 | 26,125 |
| SAT021 | MM27425 | SAT031 | 0,40 | 8,65 | 2,30E-16 | 0,11 | 192 | 174,25 | 74,5 |
| SAT031 | SAT012 | SAT021 | 0,40 | 4,80 | 1,57E-06 | 0,13 | 677,75 | 67,75 | 29 |
| SAT021 | SAT003 | TLMF Lep 30362 | 0,40 | 7,46 | 8,63E-14 | 0,09 | 224,75 | 143,5 | 61,5 |
| TLMF Lep 30361 | SAT012 | SAT003 | 0,40 | 4,08 | 4,42E-05 | 0,11 | 576,25 | 70 | 30 |

## Discussion

*S. pavonia* and *S. pavoniella* represent a well-known system of Lepidoptera with parapatric relationships having a narrow contact zone in Europe, similar to *Melitaea athalia*/*celadussa* (Joshi et al., 2022; Tahami et al., 2021). Using target enrichment approach, we used this species pair as a model to elucidate the evolutionary genetic patterns and admixture as well as species delimitation under parapatric mode of distribution. In the light of results obtained from hundreds of loci, we attempt to interpret the observations from evolutionary and taxonomic points of view, taking into consideration the past climatic oscillations that have shaped the genetic diversity of European Lepidoptera (Hewitt, 1996, 2000, 2004). Systems like the one studied here are likely to be common in nature, and while many cases are documented, many more remain undetected because parapatric taxa usually represent recent divergence of lineages at the interface of species and population levels, rendering observing the presence of two taxonomically distinct but possibly cryptic entities challenging. We find it possible that many closely related species have undergone a similar phase of a secondary contact following partial or full reproductive incompatibility evolved in isolation, particularly during glaciation maxima when ranges were split into several isolated refugia (Schmitt, 2007). Periodic expansions of species’ ranges due to climatic oscillations can be supposed to lead to many secondary contacts of such diverged populations, which in turn will provide grounds for ecological differentiation of reproductively diverged populations through reinforcement and character displacement.

### Mitonuclear discordance

The Maximum likelihood tree for mtDNA dataset did not fully separate the two taxa, suggesting that *S. pavonia* and *S. pavoniella* may not be distinct at species level (Fig. 1). In contrast, the Maximum Likelihood tree and the ASTRAL species tree from the genomic dataset recovered the two taxa as separate entities, with hybrids being grouped together in ASTRAL tree and shown to be intermediate with regards to the wild collected specimens (Figs. 1 and 2). The causes of this mitonuclear discordance could be operational or biological (Ivanov et al., 2018). Possible biological causes include incomplete lineage sorting (ILS), introgression, horizontal gene transfer, and *Wolbachia*-mediated selective sweeps following hybridization (Bergthorsson et al., 2003; Mutanen et al., 2016; Soucy et al., 2015; Toews & Brelsford, 2012). The operational causes include misidentifications, contaminations, paralogs, NUMTs and chimeric sequences, but also inconsistencies due to inaccurate taxonomy (Funk & Omland, 2003; Mutanen et al., 2016). In this study operational causes were ruled out via thorough analysis and investigation and the results were re-analyzed to systematically eliminate operational causes as contributing factors.

While it is difficult to distinguish between incomplete lineage sorting and introgression at least based on phylogenomic analyses (Funk & Omland, 2003), some insights of whether ILS is present in the dataset or not can be obtained. Generally, in presence of ILS, individual genealogies can be discordant with each other and with the species tree (Degnan & Rosenberg, 2009; Maddison, 1997). From our results, we observed that both ASTRAL, which is statistically consistent under ILS and the concatenation-based phylogenomic trees display similar relationships between the two taxa, with normalized quartet score from ASTRAL calculated as 0.53. This score is indicator of amount of gene tree discordance due to ILS, but in some cases can also suggest gene tree error (Mirarab, 2019). Thus, ILS is likely present in this dataset, but in moderate levels only.

There is a strong evidence for mitonuclear discordance taking place due to historical introgression (Linnen & Farrell, 2007; Shaw, 2002; Sota & Vogler, 2001). From STRUCTURE and geographic distribution of genomic admixture, it can be observed that the two taxa do not display significant levels of admixture. However, the two taxa are suggested to have been involved in sporadic introgression in the past caused by hybrid males (Huemer & Nässig, 2003b). Analysis of ancient introgression using Dsuite supports this idea of episodes of past introgression. However, it appears that presently the two taxa show none or only little gene flow between them.

Although the topologies of barcode and ASTRAL tree differed from each other, in both the trees we observed two specimens on Italy placed on long internal branch. In ASTRAL, such long branches are generally indicative of gene tree discordance, as the branch lengths are in coalescent units (Sayyari & Mirarab, 2016). In some cases, they could also be the indicative of presence of pseudogenes, or bad alignments due to missing data or shorter sequences, etc. As we had carefully checked all our alignments (barcode and nuclear genes) before proceeding for further analysis, the possibility of misalignments can be ruled out in both the cases and the biological causes, if any, remain to be discovered.

Results from species delimitation analyses using BPP were found to be contradictory to those suggested from all the other analyses, where *S. pavonia* and *S. pavoniella* are clearly identified as separate species. The gdi values have been shown to be unreliable predictors of the species status (Dietz et al., 2022), and gdi values estimated for taxonomically distinct species may sometimes be lower than those estimated for intraspecific entities (Jackson et al., 2017), which was observed to be the case in the present study.

### Character displacement and parapatry

Results from analysis of nuclear markers strongly suggest the two taxa are separate species and both fulfill the criteria of the biological species concept (reproductive barrier present) as well as the phylogenetic species concept (reciprocal monophyly). The apparent lack of gene flow suggests that the taxa have developed barriers to gene flow, despite having a narrow geographical overlap. This is also evident from isolation by distance (IBD) analysis, where geographic distance was not found to be significant in explaining the observed genetic separation of the two taxa. Often, species that are a part of parapatric distribution show striking similarities morphologically and ecologically, in addition to that individuals that are morphologically intermediate may occur (Guiller et al., 2017; Slender et al., 2017; Tahami et al., 2021). In the present study system, *S. pavonia* and *S. pavoniella* are morphologically similar, with *S. pavoniella* being on average larger in size. The difference between the wing patterns and the genitalia were noticed only 20 years ago, and quite some time even after being separated by Huemer & Nässig (2003b), they were commonly identified as a single species. There is a possibility of hybridization if both species are brought in close approximation since our hybrids came from breeding between the two species under laboratory conditions. But they also hybridize naturally as reported by Huemer & Nässig (2003b). Although hybridization occurs, it seems that these hybrids are sterile in natural populations (Huemer & Nässig, 2003b).

Two possible scenarios commonly emerge while attempting to explain the evolutionary dynamics of closely related parapatric taxa; parapatric speciation, or secondary contact of the populations diverged in allopatry due to range expansions. In the present study system, the interspecific interaction in the zone of overlap could have generated either ecological or reproductive character displacement. There is evidence that ecological character displacement can promote speciation as the diverging sub-populations are adapted to different habitats and eventually attain reproductive isolation (Reifová et al., 2011). The two species share very much the same ecological niche and are attracted to the pheromones of the same species, suggesting that they have not been in contact long time as pheromonal differentiation would then be expected in species like *Saturnia* where pheromones play a critical role in mate attraction. This is inconsistent with the observation of ecological separation (due to adaptation to local habitats) during parapatric speciation (Coyne & Orr, 2004).

Although we cannot fully rule out the possibility of parapatric speciation, presumably, the two *Saturnia* populations were separated during the last interglacial period in distinct refugia and differentiated in local allopatry due to appearance of prezygotic isolation factors, followed by the post-glacial expansion of their ranges resulting in secondary contact, as has been assumed for other species (Tahami et al., 2021). Although we cannot exactly know whether these pre-zygotic isolation factors arose in allopatry or during the secondary contact of the diverged populations. It is known that the body size is sometimes a potential target for selection under reproductive character displacement when two species come into contact, as body size is closely associated with the mating behavior (Zhang & Kubota, 2023). In theory, the secondary contact of diverged populations may either result in speciation through reinforcement (Pfennig, 2016), or the merging of the populations. In the present study system, species-level differentiation between the *S. pavonia* and *S. pavoniella* as indicated by most of the analyses suggest that the second possibility was unlikely to happen, as there is a mechanism at play that is maintaining a genetic barrier between two taxa. There is evidence of past introgression as Dsuite results suggest, but it has not yet resulted in disruption of this barrier. In some cases, hybrid zones can also act as a ‘genetic sink’, that strongly restricts gene flow between the species (Hafner et al., 1983; Yanchukov et al., 2006). Indeed, it has been suggested that the divergence in local allopatry is protected by hybrid zones as the taxa later become parapatric (Hewitt 1989). However, to test this idea in the present study system, a more detailed study of samples from the contact zone would be required.

Parapatric species systems provide interesting models to study speciation in action. Study of such systems take place at the interface of population genetics and phylogenetics levels for which reason analyses designed to address question at both levels are potentially useful. Studied cases of parapatric species pairs or groups have commonly revealed patterns that suggest speciation not having reached completion, making taxonomic delimitation of parapatric taxa often very challenging and largely arbitrary, and highlighting the nature of speciation as a slow gradual process rather than an event. From an evolutionary point of view, one might expect observing systems at different stages of the speciation-population continuum, particularly when parapatry has resulted from a secondary contact of populations differentiated in allopatry, for example during glacial refugial periods.

### Potential of target enrichment in species delimitation

Studies in the last decade have suggested and favored the use of multi-locus genomic markers such as ultraconserved elements (UCEs), ddRAD-sequencing and target capture for species delimitation (Natusch et al., 2020; Rancilhac et al., 2019; Smith et al., 2014). In the present study, we used LepZFMK 1.0 kit developed by Mayer et al. (2021), which targets 2953 CDS regions in 1753 nuclear genes. This approach was adopted from Joshi et al. (2022) that reported a successful capture experiment and concluded that loci used in a target enrichment experiment, particularly with flanking regions included, contain sufficient phylogenetic information for the delimitation of closely related species. The number of loci recovered from our genomic dataset were found to be sufficient to infer the phylogenetic relationships between *S. pavonia* and *S. pavoniella* as can be seen from our phylogenomic analyses. The percentage of missing data ranged from 18.56% to 35.08%, which is modest compared for example to those typically seen in RAD methods between species (Lee et al., 2018). It is an inexpensive and cost-effective approach as compared to whole genome sequencing (WGS) and whole exome sequencing (WES) with greater sequencing depth and less data burden (Bewicke-Copley et al., 2019). Banker et al., (2020) reports that target enrichment methods helps researchers to choose genetic markers at both deep- and shallow-scales of taxonomy that further facilitates in inferring phylogenetic relationships among closely related taxa. Other alternatives to target enrichment for this study could have been RAD-seq, which could not possibly match the efficiency of target enrichment due to their limitations such as locus-dropout effect, high percentage of missing data, etc. (Banker et al., 2020; Lee et al., 2018).

## Conclusions

In this study, we utilized the Lepidoptera system of *S. pavonia* and *S. pavoniella* to examine evolutionary genetic patterns, species delimitation, and admixture in a parapatric mode of distribution. Our findings support the hypothesis that the two taxa are separate species based on both biological and phylogenetic species concepts. However, there is evidence of mitonuclear discordance due to past introgression, and incomplete lineage sorting may also be present. The study also suggests that many closely related species may have experienced a similar phase of secondary contact after isolation, and periodic range expansions due to climatic oscillations may result in many secondary contacts of such diverged populations, which may lead to ecological differentiation. Our study highlights the importance of understanding the evolutionary processes that shape the genetic diversity of organisms in parapatric relationships, which are likely to be common in nature. We found that the taxonomy of parapatric species can sometimes be straightforward, as we could not detect ongoing hybridization and introgression in *Saturnia*. However, our results and observations suggest that the two reproductively isolated taxa have not yet completed speciation from an ecological standpoint, as their sexual pheromones continue to attract heterospecific individuals and hybridizations occur. We propose that the lack of pheromonal differentiation plays a significant evolutionary role in maintaining parapatry, as individuals dispersing into the area of the other species are likely to have low fitness. This situation creates evolutionary “pressure” for the two species to develop more reliable prezygotic isolation mechanisms through pheromonal character displacement. Although we lack direct evidence, we hypothesize that this process is underway. Conducting detailed studies focusing on the natural pheromone chemistry of the two species in and around the contact zone would provide valuable insights into this process.

## Acknowledgements

This study was supported by the Academy of Finland through Grant No. 314702 to MM. CLV was partly funded by FEDER project InfoBioS (EX011185). MJ would like to thank Finnish cultural foundation and University of Oulu scholarship foundation for providing funding to continue her doctoral studies. We would like to thank Laura Törmälä for her assistance in laboratory procedures and Vladislav Ivanov for constructive discussions regarding Dsuite analyses. Thanks are due to Till Tolasch for synthesizing and sending us the sex pheromone that was used to attract and collect specimens for this study. We thank Carlos Antonietty Adame, Yann Baillet, Philippe Bordet, Gregory Guicherd, Stanislav Gomboc, Zdenek Lastuvka and Enrique Murria Beltran for sending *Saturnia* specimens from their respective home countries. The sequencing service was provided by the Biomedicum Functional Genomics Unit at the Helsinki Institute of Life Science and Biocenter Finland at the University of Helsinki. Additionally, we would like to thank CSC Finland for providing computational resources to carry out bioinformatics analyses.

## Data Accessibility

The resulting sequences, along with the voucher data and images, are deposited in the Barcode of Life Database (BOLD), http://www.boldsystems.org (Ratnasingham and Hebert, 2007) and the sequences subsequently deposited in GenBank. All data are available from BOLD in the dataset DS-PAVONI (dx.doi.org/XX/DS-PAVONI).

Raw target enrichment sequencing data is archived on the NCBI Sequence Read Archive under BioProject ID PRJNA933196. Alignments, treefiles and other data is to be uploaded to Zenodo repository.

## Benefit Sharing Statement

Not Applicable

## Author Contributions

MM conceived the presented idea. CLV and PH provided the materials for the study. ME generated the target enrichment bait kit. MJ designed the research. MK and MJ performed the experiments, analyzed the data as well as wrote the paper. All authors discussed the results and contributed to the final manuscript.

## Notes

### Competing Interest Statement

The authors have declared no competing interest.

