## Supplementary_File for "Patterns of speciation in a parapatric pair of *Saturnia* moths as revealed by Target Capture"

**Supplementary information for manuscript ‘Patterns of speciation in a parapatric pair of *Saturnia* moths as revealed by Target Capture’**

^1^ Maria Khan*, ^1^ Mukta Joshi*, ^2^ Marianne Espeland, ^3^ Peter Huemer, ^4,5^ Carlos Lopez Vaamonde, ^1^ Marko Mutanen

**Barcoding laboratory protocol –**

The COI was amplified for using the primers HybLCO and HybHCO, and PCR was conducted under the following conditions: 95c 5min, (40x 95c 30s, 50c 30s, 72c 2min) and 72c 2min.

PCR purification was done using the Exo-sap purification method according to the following instructions - PCR product 4µl, Fast AP 0.5µl, FA buffer 0.8, EXO I 0.1µl and water 4.6µl. The final volume of the purified product was 10µl.

5µl of purified PCR product and 5µl of 5µm primer were sent for sequencing.

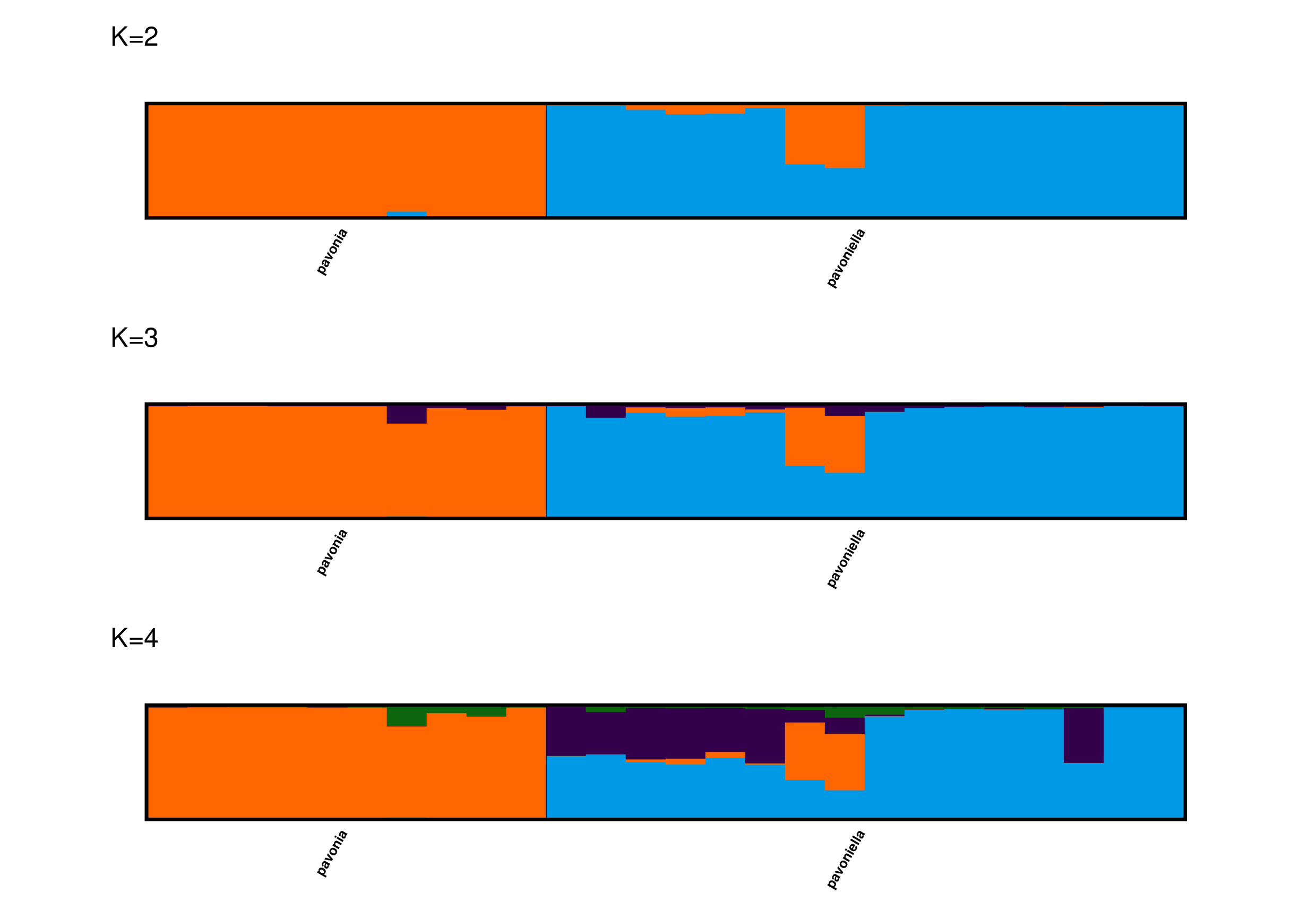

**Figure S1**: STRUCTURE barplots obtained using CLUMPAK program showing cluster assignments from K=2 toK=4.

a)

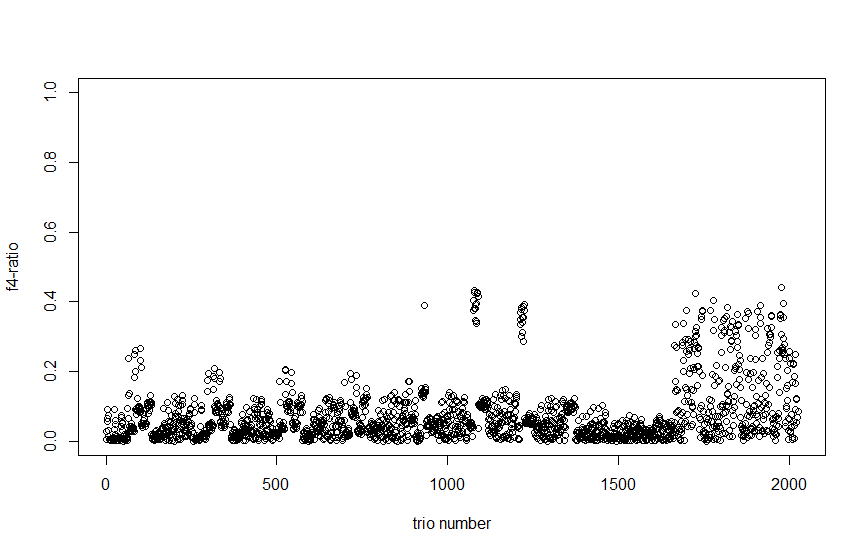

b)

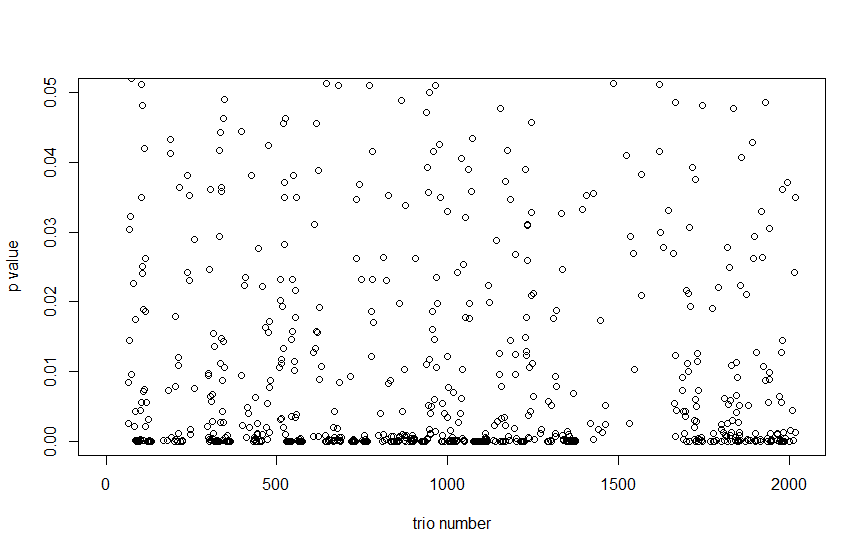

c)

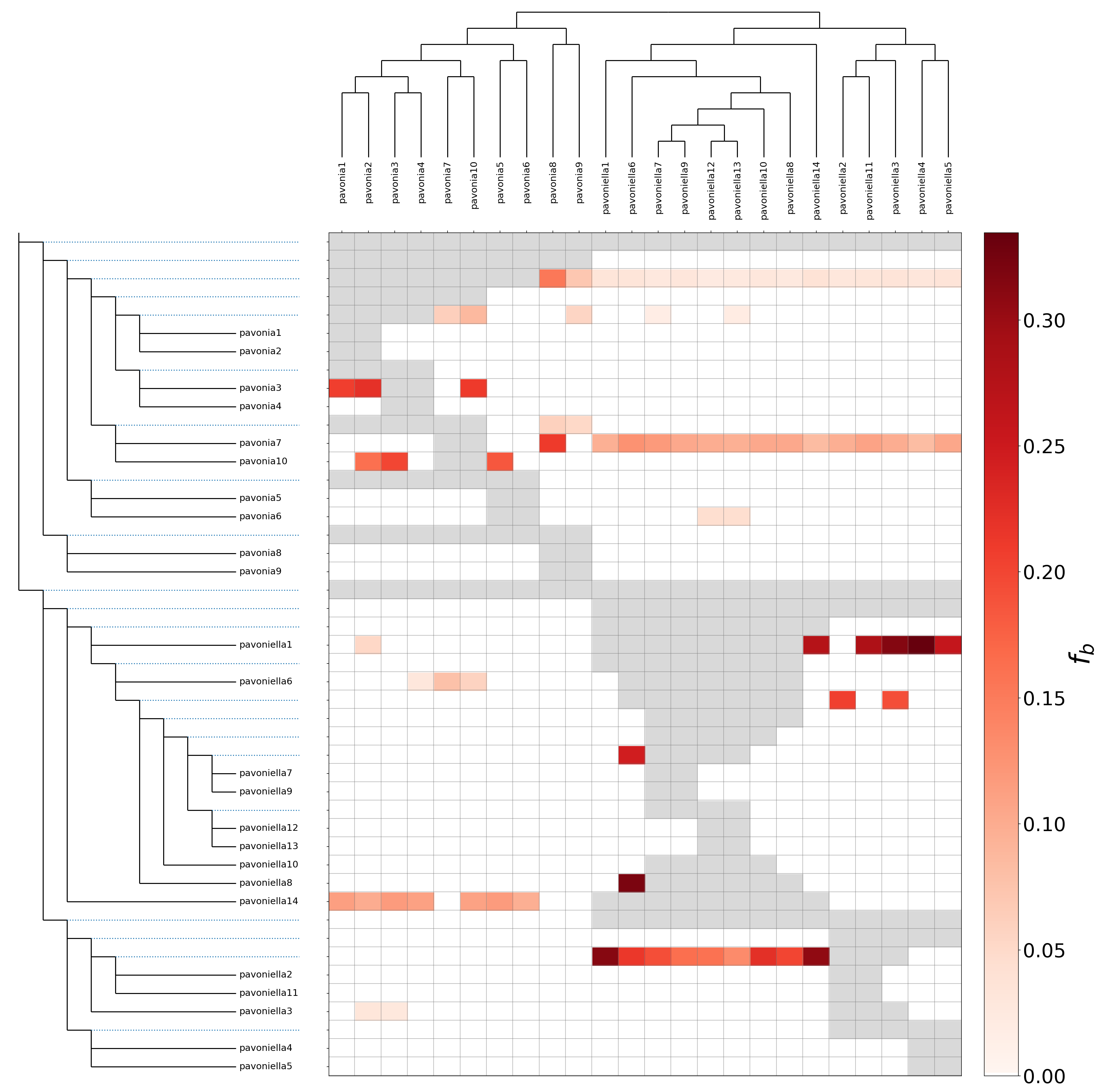

**Figure S2**: a) f4-ratio, or proportion of the genome affected by gene flow for all the trios b) BH-corrected p-value (Benjamini-Hochberg) for all the trios c) The introgression proportions mapped to the internal branches of ASTRAL species tree using *f*-branch approach. The gray cells represent pairs of species or branches that cannot be tested (individuals and corresponding sample codes - pavonia1: SAT003, pavonia2: SAT004, pavoniella1: SAT006, pavonia3: SAT008, pavoniella2: SAT009, pavoniella3: SAT010, pavoniella4: SAT011, pavoniella5: SAT012, pavoniella14: SAT013, pavonia4: CLV8286, pavonia5: CLV8287, pavonia6: CLV8290, pavonia7: CLV8292, pavoniella6: CLV8294, pavoniella7: CLV8297, pavoniella8: CLV8299, pavonia8: MM27425, pavonia9: MM27428, pavoniella9: TLMF Lep 03073, pavonia10: TLMF Lep 30352, pavoniella10: TLMF Lep 30357, pavoniella11: TLMF Lep 30358, pavoniella12: TLMF Lep 30361, pavoniella13: TLMF Lep 30362)

a)

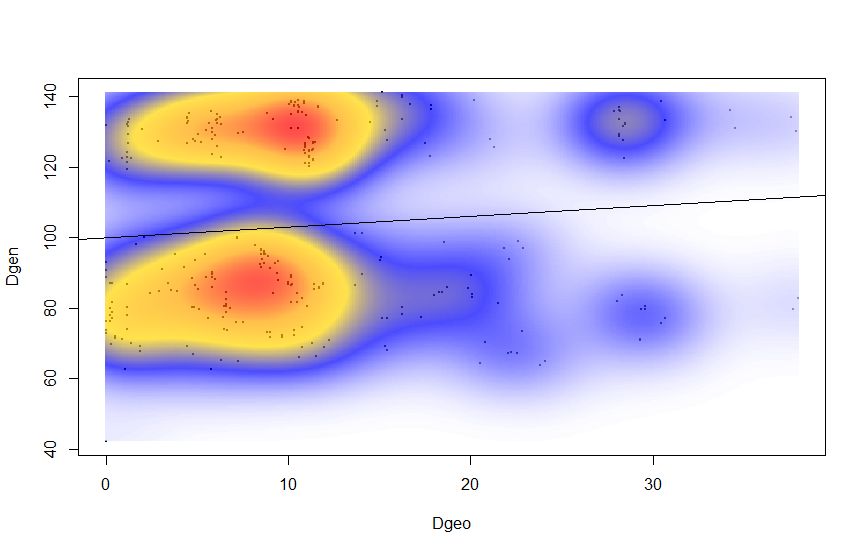

b)

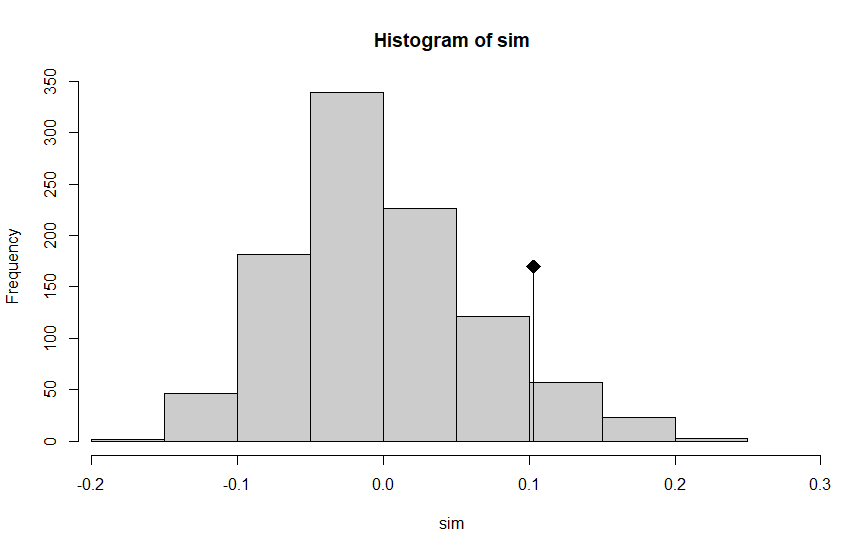

P-value= 0.075

**Figure S3**: a) The scatterplot of the correlation matrix, where local density is measured using 2-dimentional kernel density estimation b) The histograms of permuted values (in absence of a spatial structure) of a correlation between two distance matrices – Edwards’ distance and Euclidean geographic distance. The original value of a correlation is represented by the dot, which represents a significant spatial structure if it is out of the reference distribution

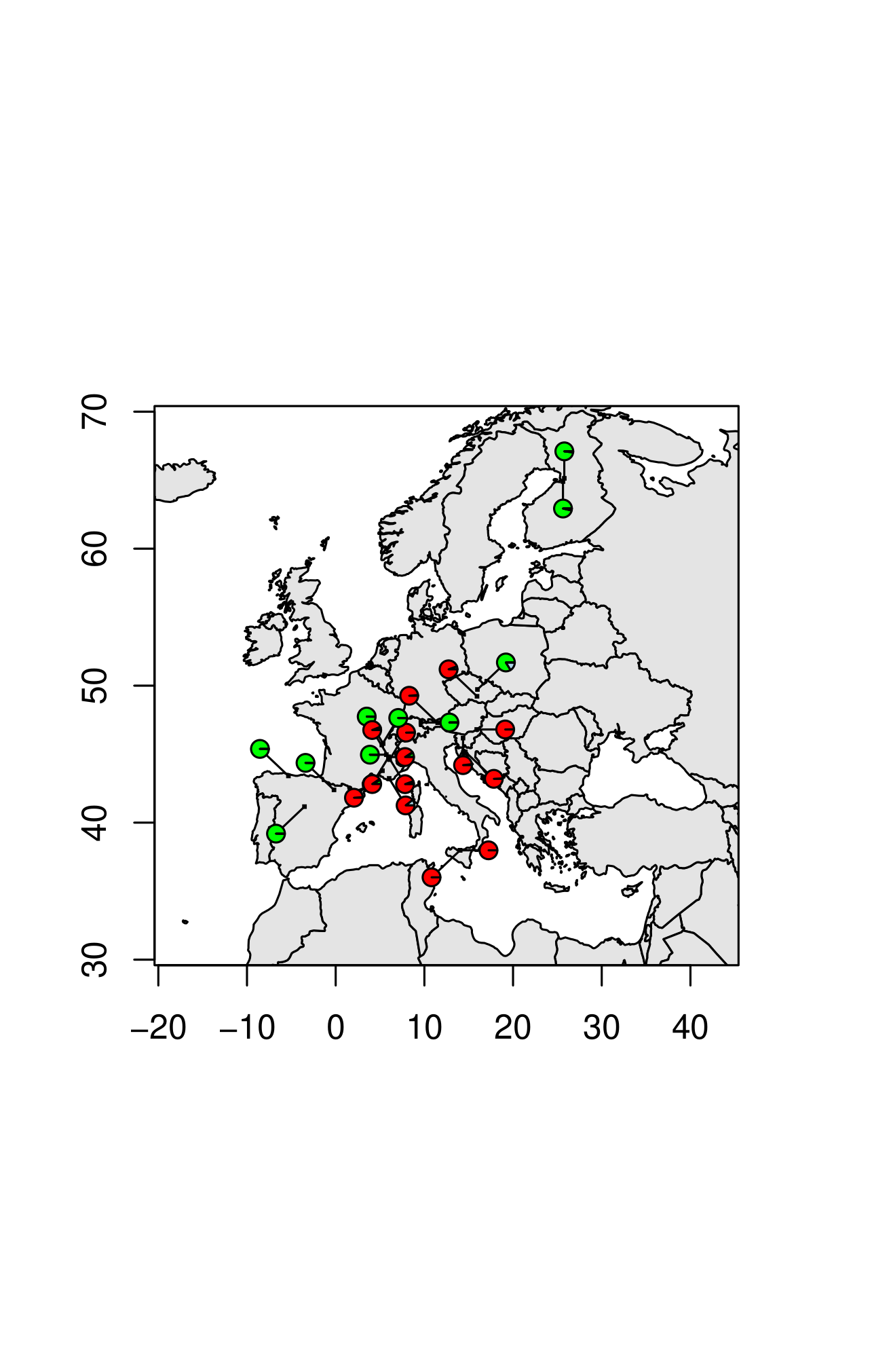

**Figure S4**: Pie charts based on membership coefficient matrix at K=3 mapped on geographic coordinates

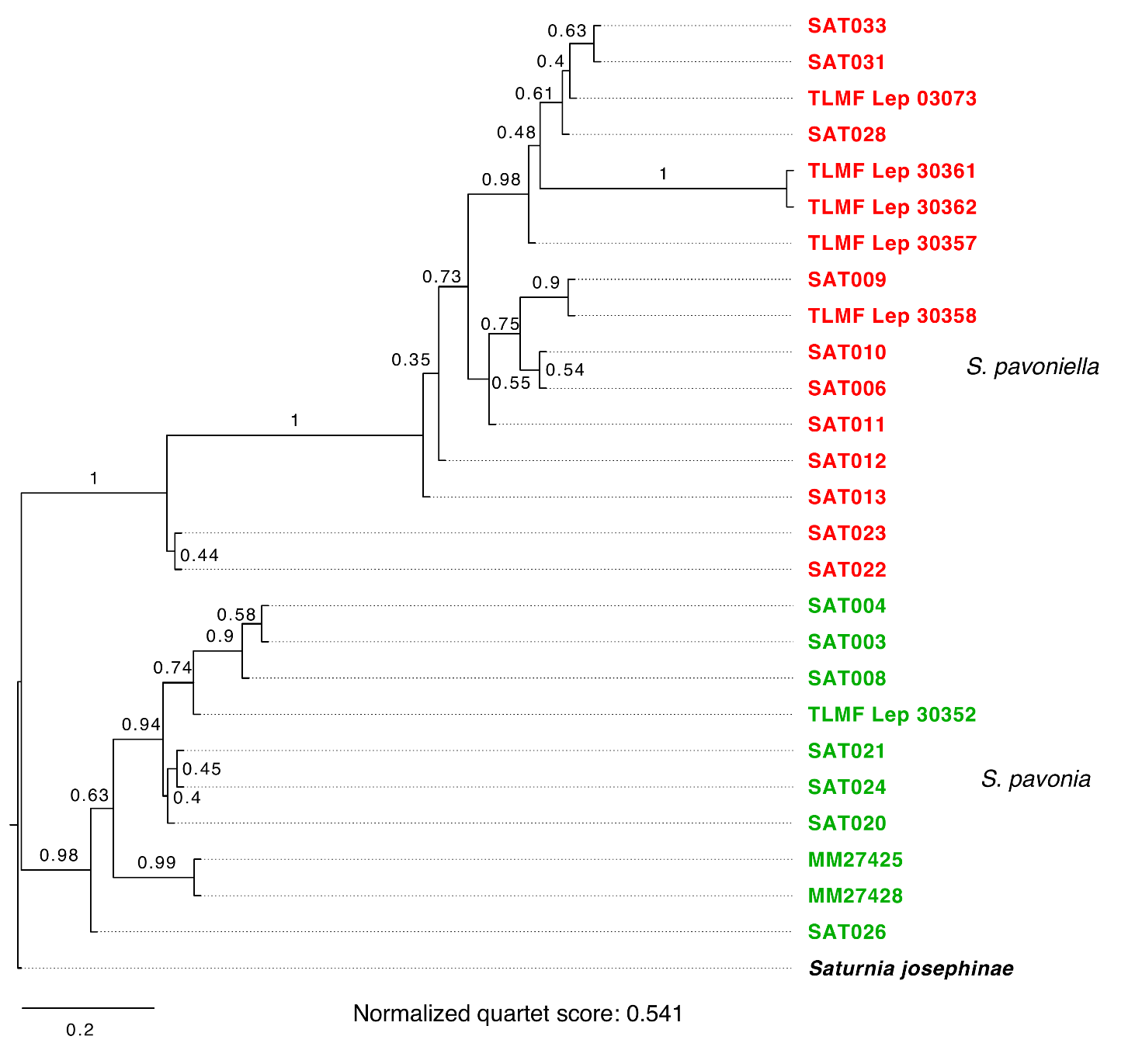

**Figure S5**: ASTRAL tree obtained from dataset filtered using Treeshrink

**Table S1**. Delta K statistics calculated for each cluster based on the SNP dataset using Evanno method (Evanno et al., 2005) for STRUCTURE run.

| K | Reps | Mean LnP(K) | | Stdev LnP(K) | Ln'(K) | \|Ln''(K)\| | Delta K |
| --- | --- | --- | --- | --- | --- | --- | --- |
| 1 | 10 | -16581.46 | | 7.246 | NA | NA | NA |
| 2 | 10 | -12752.62 | | 2.0746 | 3828.84 | 4023.51 | 1939.406977 |
| 3 | 10 | -12947.29 | | 20.806 | -194.67 | 5496.84 | 264.194867 |
| 4 | 10 | -18638.8 | | 9927.0575 | -5691.51 | 4472.94 | 0.450581 |
| 5 | 10 | -19857.37 | | 11775.0477 | -1218.57 | NA | NA |

**Table S2:** D-statistics values > 0.25 with p-value < 0.05 (individuals and corresponding sample codes - pavonia1: SAT003, pavonia2: SAT004, pavoniella1: SAT006, pavonia3: SAT008, pavoniella2: SAT009, pavoniella3: SAT010, pavoniella4: SAT011, pavoniella5: SAT012, pavoniella14: SAT013, pavonia4: CLV8286, pavonia5: CLV8287, pavonia6: CLV8290, pavonia7: CLV8292, pavoniella6: CLV8294, pavoniella7: CLV8297, pavoniella8: CLV8299, pavonia8: MM27425, pavonia9: MM27428, pavoniella9: TLMF Lep 03073, pavonia10: TLMF Lep 30352, pavoniella10: TLMF Lep 30357, pavoniella11: TLMF Lep 30358, pavoniella12: TLMF Lep 30361, pavoniella13: TLMF Lep 30362)

| **P1** | **P2** | **P3** | **Dstatistic** | **Z-score** | **p-value** | **f4-ratio** | **BBAA** | **ABBA** | **BABA** |
| --- | --- | --- | --- | --- | --- | --- | --- | --- | --- |
| pavoniella13 | pavonia7 | pavonia5 | 0,517875 | 10,0141 | 2,30E-16 | 0,433704 | 226,75 | 185,75 | 59 |
| pavoniella12 | pavonia7 | pavonia5 | 0,493374 | 7,69528 | 1,41E-14 | 0,427562 | 219,375 | 183,125 | 62,125 |
| pavonia4 | pavonia7 | pavoniella13 | 0,49001 | 10,1871 | 2,30E-16 | 0,142383 | 213,25 | 195,75 | 67 |
| pavoniella12 | pavoniella5 | pavonia4 | 0,489247 | 8,69022 | 2,30E-16 | 0,138931 | 613,25 | 69,25 | 23,75 |
| pavoniella9 | pavonia7 | pavonia5 | 0,486052 | 6,84296 | 7,76E-12 | 0,415215 | 222,375 | 173,125 | 59,875 |
| pavonia4 | pavonia7 | pavoniella7 | 0,485359 | 9,39217 | 2,30E-16 | 0,155177 | 216,75 | 209,25 | 72,5 |
| pavonia4 | pavonia7 | pavoniella10 | 0,48503 | 9,19427 | 2,30E-16 | 0,150651 | 200,25 | 186 | 64,5 |
| pavoniella12 | pavoniella5 | pavonia5 | 0,483871 | 6,36376 | 1,97E-10 | 0,145503 | 646 | 63,25 | 22 |
| pavoniella7 | pavonia7 | pavonia5 | 0,481518 | 7,33768 | 2,17E-13 | 0,425623 | 243,875 | 190,375 | 66,625 |
| pavoniella8 | pavonia7 | pavonia5 | 0,480658 | 7,85769 | 3,91E-15 | 0,423339 | 236,625 | 191,375 | 67,125 |
| pavoniella13 | pavoniella5 | pavonia5 | 0,480447 | 6,41336 | 1,42E-10 | 0,14726 | 658,75 | 66,25 | 23,25 |
| pavonia4 | pavonia7 | pavoniella12 | 0,476654 | 9,15572 | 2,30E-16 | 0,138692 | 212,5 | 189,75 | 67,25 |
| pavoniella10 | pavonia7 | pavonia5 | 0,465859 | 7,6647 | 1,79E-14 | 0,403626 | 220,375 | 166,375 | 60,625 |
| pavonia4 | pavonia7 | pavoniella9 | 0,462598 | 8,60462 | 2,30E-16 | 0,147382 | 195,25 | 185,75 | 68,25 |
| pavonia4 | pavonia7 | pavoniella8 | 0,461608 | 9,23621 | 2,30E-16 | 0,139656 | 220,75 | 202,25 | 74,5 |
| pavoniella12 | pavoniella14 | pavonia4 | 0,457534 | 7,47446 | 7,75E-14 | 0,130469 | 566,75 | 66,5 | 24,75 |
| pavoniella6 | pavonia7 | pavonia5 | 0,45283 | 6,79563 | 1,08E-11 | 0,395242 | 242,5 | 173,25 | 65,25 |
| pavoniella6 | pavonia7 | pavonia4 | 0,451524 | 9,22291 | 2,30E-16 | 0,389021 | 208,75 | 196,5 | 74,25 |
| pavoniella13 | pavoniella14 | pavonia4 | 0,449036 | 7,11377 | 1,13E-12 | 0,127443 | 573,5 | 65,75 | 25 |
| pavoniella13 | pavoniella14 | pavonia5 | 0,443077 | 5,52501 | 3,29E-08 | 0,129964 | 615,375 | 58,625 | 22,625 |
| pavoniella13 | pavoniella5 | pavonia4 | 0,441624 | 6,82962 | 8,51E-12 | 0,131123 | 625 | 71 | 27,5 |
| pavoniella12 | pavoniella14 | pavonia5 | 0,440367 | 5,24854 | 1,53E-07 | 0,131387 | 605,625 | 58,875 | 22,875 |
| pavoniella12 | pavoniella5 | pavonia3 | 0,437956 | 5,25142 | 1,51E-07 | 0,132159 | 591,875 | 73,875 | 28,875 |
| pavoniella2 | pavonia7 | pavonia5 | 0,435798 | 5,82624 | 5,67E-09 | 0,392638 | 228,25 | 184,5 | 72,5 |
| pavoniella3 | pavonia7 | pavonia5 | 0,434783 | 6,18417 | 6,24E-10 | 0,379377 | 208,625 | 160,875 | 63,375 |
| pavoniella12 | pavoniella14 | pavonia10 | 0,433333 | 5,78161 | 7,40E-09 | 0,12211 | 528,125 | 69,875 | 27,625 |
| pavonia5 | pavonia8 | pavoniella9 | 0,432492 | 8,46094 | 2,30E-16 | 0,116741 | 176,875 | 163,125 | 64,625 |
| pavoniella9 | pavonia7 | pavonia6 | 0,430928 | 7,5828 | 3,38E-14 | 0,39102 | 193,25 | 173,5 | 69 |
| pavonia5 | pavonia8 | pavoniella13 | 0,430449 | 8,79891 | 2,30E-16 | 0,105447 | 189 | 163,25 | 65 |
| pavoniella1 | pavonia7 | pavonia5 | 0,427152 | 5,70111 | 1,19E-08 | 0,376093 | 212,875 | 161,625 | 64,875 |
| pavoniella12 | pavoniella14 | pavonia3 | 0,426768 | 6,53715 | 6,27E-11 | 0,129304 | 547,375 | 70,625 | 28,375 |
| pavonia5 | pavonia9 | pavoniella9 | 0,426454 | 5,73579 | 9,71E-09 | 0,114233 | 179,125 | 156,375 | 62,875 |
| pavoniella13 | pavoniella5 | pavonia3 | 0,426374 | 5,07754 | 3,82E-07 | 0,13867 | 604,875 | 81,125 | 32,625 |
| pavoniella13 | pavoniella14 | pavonia10 | 0,425974 | 5,86531 | 4,48E-09 | 0,120323 | 536,875 | 68,625 | 27,625 |
| pavonia3 | pavonia7 | pavoniella6 | 0,422523 | 7,10402 | 1,21E-12 | 0,15061 | 218,875 | 197,375 | 80,125 |
| pavoniella13 | pavonia7 | pavonia6 | 0,422287 | 9,09163 | 2,30E-16 | 0,383319 | 188,375 | 181,875 | 73,875 |
| pavoniella11 | pavonia7 | pavonia5 | 0,422009 | 6,1872 | 6,12E-10 | 0,380906 | 223,375 | 166,375 | 67,625 |
| pavoniella13 | pavoniella6 | pavonia4 | 0,420408 | 3,11722 | 0,001826 | 0,076808 | 685,25 | 43,5 | 17,75 |
| pavonia4 | pavonia7 | pavoniella2 | 0,419503 | 9,3961 | 2,30E-16 | 0,132327 | 216,875 | 192,875 | 78,875 |
| pavoniella12 | pavoniella14 | pavonia1 | 0,418824 | 5,30917 | 1,10E-07 | 0,128058 | 532,875 | 75,375 | 30,875 |
| pavoniella8 | pavoniella5 | pavonia5 | 0,418079 | 4,81348 | 1,48E-06 | 0,127257 | 694,5 | 62,75 | 25,75 |
| pavoniella8 | pavoniella5 | pavonia4 | 0,41791 | 6,35467 | 2,09E-10 | 0,125 | 663 | 71,25 | 29,25 |
| pavoniella13 | pavoniella11 | pavonia5 | 0,41791 | 3,40654 | 0,000658 | 0,096719 | 703,25 | 47,5 | 19,5 |
| pavoniella9 | pavoniella14 | pavonia5 | 0,416918 | 4,59321 | 4,36E-06 | 0,130066 | 603,125 | 58,625 | 24,125 |
| pavonia5 | pavonia8 | pavoniella6 | 0,41608 | 10,6988 | 2,30E-16 | 0,122885 | 179,625 | 176,125 | 72,625 |
| pavoniella9 | pavoniella14 | pavonia1 | 0,416076 | 5,25459 | 1,48E-07 | 0,130081 | 524,125 | 74,875 | 30,875 |
| pavoniella8 | pavoniella5 | pavonia3 | 0,414474 | 6,08166 | 1,19E-09 | 0,132446 | 640,625 | 80,625 | 33,375 |
| pavoniella7 | pavonia7 | pavonia6 | 0,412959 | 7,48967 | 6,90E-14 | 0,386775 | 205,875 | 182,625 | 75,875 |
| pavonia5 | pavonia9 | pavoniella13 | 0,41217 | 5,25462 | 1,48E-07 | 0,099778 | 195,25 | 153,75 | 64 |
| pavoniella14 | pavoniella2 | pavoniella6 | 0,411067 | 6,13715 | 8,40E-10 | 0,29686 | 142,125 | 133,875 | 55,875 |
| pavoniella12 | pavoniella11 | pavonia4 | 0,410646 | 4,99204 | 5,97E-07 | 0,081387 | 659,875 | 46,375 | 19,375 |
| pavonia4 | pavonia7 | pavoniella11 | 0,409821 | 9,68391 | 2,30E-16 | 0,133415 | 195,125 | 186,625 | 78,125 |
| pavoniella12 | pavoniella5 | pavonia2 | 0,409548 | 5,20768 | 1,91E-07 | 0,122281 | 566,875 | 70,125 | 29,375 |
| pavonia5 | pavonia8 | pavoniella12 | 0,408186 | 7,0788 | 1,45E-12 | 0,101179 | 185,875 | 159,125 | 66,875 |
| pavoniella12 | pavoniella4 | pavonia2 | 0,407311 | 4,99232 | 5,97E-07 | 0,120092 | 530,125 | 67,375 | 28,375 |
| pavonia3 | pavonia7 | pavoniella7 | 0,40724 | 7,05243 | 1,76E-12 | 0,13365 | 237,125 | 194,375 | 81,875 |
| pavonia5 | pavonia9 | pavoniella6 | 0,406417 | 5,8447 | 5,07E-09 | 0,11827 | 186,125 | 164,375 | 69,375 |
| pavonia5 | pavonia2 | pavoniella9 | 0,40613 | 5,94364 | 2,79E-09 | 0,096804 | 204,375 | 137,625 | 58,125 |
| pavoniella12 | pavoniella14 | pavonia2 | 0,405914 | 5,53933 | 3,04E-08 | 0,116873 | 528,625 | 65,375 | 27,625 |
| pavoniella12 | pavonia7 | pavonia6 | 0,404878 | 9,44668 | 2,30E-16 | 0,37693 | 188,75 | 180 | 76,25 |
| pavonia5 | pavonia2 | pavoniella6 | 0,403571 | 7,20585 | 5,77E-13 | 0,104986 | 200,125 | 147,375 | 62,625 |
| pavoniella10 | pavonia7 | pavonia6 | 0,403545 | 7,24804 | 4,23E-13 | 0,370335 | 189 | 168,25 | 71,5 |
| pavonia2 | pavonia7 | pavoniella7 | 0,40321 | 6,39789 | 1,58E-10 | 0,122486 | 226,125 | 174,875 | 74,375 |
| pavoniella13 | pavoniella14 | pavonia3 | 0,402632 | 6,13231 | 8,66E-10 | 0,116527 | 554,125 | 66,625 | 28,375 |
| pavonia5 | pavonia9 | pavoniella10 | 0,401766 | 5,35775 | 8,43E-08 | 0,109837 | 175,5 | 158,75 | 67,75 |
| pavoniella10 | pavoniella5 | pavonia5 | 0,401146 | 5,3375 | 9,42E-08 | 0,134357 | 644,125 | 61,125 | 26,125 |
| pavonia5 | pavonia8 | pavoniella7 | 0,401005 | 8,65041 | 2,30E-16 | 0,109435 | 192 | 174,25 | 74,5 |
| pavoniella7 | pavoniella5 | pavonia5 | 0,400517 | 4,80167 | 1,57E-06 | 0,133851 | 677,75 | 67,75 | 29 |
| pavonia5 | pavonia1 | pavoniella13 | 0,4 | 7,46041 | 8,63E-14 | 0,088865 | 224,75 | 143,5 | 61,5 |
| pavoniella12 | pavoniella5 | pavonia1 | 0,4 | 4,08432 | 4,42E-05 | 0,108181 | 576,25 | 70 | 30 |
| pavonia4 | pavonia7 | pavoniella1 | 0,399803 | 7,29028 | 3,09E-13 | 0,136456 | 190,25 | 177,25 | 76 |
| pavoniella13 | pavoniella14 | pavonia1 | 0,399522 | 5,43427 | 5,50E-08 | 0,119885 | 540,625 | 73,125 | 31,375 |
| pavonia5 | pavonia2 | pavoniella3 | 0,399519 | 6,47825 | 9,28E-11 | 0,11405 | 190,375 | 145,375 | 62,375 |
| pavonia5 | pavonia2 | pavoniella1 | 0,399516 | 6,2464 | 4,20E-10 | 0,107212 | 188,5 | 144,5 | 62 |
| pavoniella8 | pavonia7 | pavonia6 | 0,399247 | 9,07391 | 2,30E-16 | 0,373898 | 195,25 | 185,75 | 79,75 |
| pavoniella12 | pavoniella6 | pavonia4 | 0,398406 | 3,58132 | 0,000342 | 0,074963 | 665,125 | 43,875 | 18,875 |
| pavonia3 | pavonia7 | pavoniella13 | 0,39823 | 6,04864 | 1,46E-09 | 0,117733 | 230,5 | 177,75 | 76,5 |
| pavonia5 | pavonia10 | pavoniella9 | 0,397561 | 5,90161 | 3,60E-09 | 0,098489 | 205,75 | 143,25 | 61,75 |
| pavonia5 | pavonia10 | pavoniella13 | 0,397196 | 7,02304 | 2,17E-12 | 0,092191 | 218 | 149,5 | 64,5 |
| pavonia3 | pavonia7 | pavoniella10 | 0,395727 | 6,58471 | 4,56E-11 | 0,125443 | 212 | 171,5 | 74,25 |
| pavoniella13 | pavoniella5 | pavonia2 | 0,394009 | 4,67286 | 2,97E-06 | 0,125275 | 579,625 | 75,625 | 32,875 |
| pavonia5 | pavonia1 | pavoniella14 | 0,393692 | 6,37269 | 1,86E-10 | 0,122635 | 185,375 | 149,125 | 64,875 |
| pavonia4 | pavonia7 | pavoniella3 | 0,392644 | 9,3715 | 2,30E-16 | 0,139428 | 186,125 | 175,125 | 76,375 |
| pavonia5 | pavonia8 | pavoniella10 | 0,392585 | 8,21184 | 2,30E-16 | 0,10551 | 175,125 | 159,625 | 69,625 |
| pavonia5 | pavonia2 | pavoniella10 | 0,392252 | 6,18275 | 6,30E-10 | 0,098182 | 196,25 | 143,75 | 62,75 |
| pavoniella8 | pavoniella5 | pavonia2 | 0,392252 | 5,04498 | 4,54E-07 | 0,117818 | 609,625 | 71,875 | 31,375 |
| pavoniella8 | pavoniella14 | pavonia10 | 0,392157 | 5,56168 | 2,67E-08 | 0,113879 | 557,75 | 71 | 31 |
| pavoniella6 | pavoniella2 | pavoniella4 | 0,391778 | 5,62868 | 1,82E-08 | 0,43962 | 150,625 | 143,875 | 62,875 |
| pavonia2 | pavonia7 | pavoniella6 | 0,391304 | 6,97628 | 3,03E-12 | 0,131053 | 205 | 172 | 75,25 |
| pavoniella7 | pavoniella14 | pavonia10 | 0,391101 | 5,34645 | 8,97E-08 | 0,11844 | 555 | 74,25 | 32,5 |
| pavonia3 | pavonia7 | pavoniella8 | 0,390783 | 5,6989 | 1,21E-08 | 0,122296 | 244,125 | 188,625 | 82,625 |
| pavonia5 | pavonia8 | pavoniella8 | 0,390707 | 8,27509 | 2,30E-16 | 0,098013 | 197,125 | 164,625 | 72,125 |
| pavoniella10 | pavoniella5 | pavonia4 | 0,390374 | 6,04534 | 1,49E-09 | 0,11478 | 605,75 | 65 | 28,5 |
| pavonia10 | pavonia7 | pavoniella7 | 0,389222 | 7,0998 | 1,25E-12 | 0,118721 | 245 | 174 | 76,5 |
| pavoniella8 | pavoniella14 | pavonia3 | 0,387654 | 6,87848 | 6,05E-12 | 0,117779 | 577 | 70,25 | 31 |
| pavonia5 | pavonia9 | pavoniella7 | 0,387234 | 5,14566 | 2,67E-07 | 0,104179 | 201,5 | 163 | 72 |
| pavonia3 | pavonia7 | pavoniella9 | 0,387065 | 5,49499 | 3,91E-08 | 0,125403 | 218,5 | 174,25 | 77 |
| pavonia1 | pavonia7 | pavoniella6 | 0,386876 | 7,49617 | 6,57E-14 | 0,130795 | 218,75 | 177 | 78,25 |
| pavoniella13 | pavoniella6 | pavonia8 | 0,38676 | 3,44743 | 0,000566 | 0,074899 | 635,25 | 49,75 | 22 |
| pavonia5 | pavonia9 | pavoniella12 | 0,384966 | 4,82823 | 1,38E-06 | 0,095346 | 192,5 | 152 | 67,5 |
| pavoniella9 | pavoniella5 | pavonia5 | 0,384393 | 4,00076 | 6,31E-05 | 0,121572 | 633,875 | 59,875 | 26,625 |
| pavoniella6 | pavonia7 | pavonia6 | 0,383984 | 7,32236 | 2,44E-13 | 0,354502 | 207,75 | 168,5 | 75 |
| pavoniella8 | pavoniella14 | pavonia5 | 0,383234 | 4,46296 | 8,08E-06 | 0,114798 | 636,75 | 57,75 | 25,75 |
| pavoniella8 | pavoniella14 | pavonia4 | 0,383202 | 6,33688 | 2,34E-10 | 0,112828 | 591,375 | 65,875 | 29,375 |
| pavoniella9 | pavoniella14 | pavonia3 | 0,382653 | 4,64932 | 3,33E-06 | 0,118671 | 539,5 | 67,75 | 30,25 |
| pavonia5 | pavonia2 | pavoniella13 | 0,382426 | 5,78647 | 7,19E-09 | 0,085454 | 216,375 | 139,625 | 62,375 |
| pavonia5 | pavonia1 | pavoniella6 | 0,382181 | 6,68775 | 2,27E-11 | 0,098638 | 212,375 | 147,375 | 65,875 |
| pavoniella8 | pavoniella14 | pavonia1 | 0,381395 | 4,78569 | 1,70E-06 | 0,116065 | 556,75 | 74,25 | 33,25 |
| pavonia5 | pavonia9 | pavoniella3 | 0,381215 | 5,0688 | 4,00E-07 | 0,11946 | 170,25 | 156,25 | 70 |
| pavonia2 | pavonia7 | pavoniella9 | 0,380688 | 6,43755 | 1,21E-10 | 0,114333 | 207,75 | 155,5 | 69,75 |
| pavonia5 | pavonia10 | pavoniella10 | 0,380675 | 6,83178 | 8,39E-12 | 0,097934 | 198,5 | 148,25 | 66,5 |
| pavoniella13 | pavoniella4 | pavonia2 | 0,380247 | 4,4691 | 7,85E-06 | 0,116755 | 538,125 | 69,875 | 31,375 |
| pavonia3 | pavonia7 | pavoniella12 | 0,379921 | 6,26258 | 3,79E-10 | 0,114303 | 233,25 | 175,25 | 78,75 |
| pavoniella13 | pavoniella14 | pavonia2 | 0,379032 | 5,59791 | 2,17E-08 | 0,108712 | 536,875 | 64,125 | 28,875 |
| pavoniella2 | pavoniella5 | pavonia2 | 0,378713 | 3,91557 | 9,02E-05 | 0,111273 | 656,625 | 69,625 | 31,375 |
| pavoniella11 | pavonia7 | pavonia6 | 0,37871 | 6,65656 | 2,80E-11 | 0,348399 | 184,625 | 168,375 | 75,875 |
| pavoniella2 | pavonia7 | pavonia6 | 0,378456 | 7,98523 | 1,40E-15 | 0,356054 | 187,75 | 180,75 | 81,5 |
| pavoniella12 | pavoniella4 | pavonia4 | 0,378016 | 4,89803 | 9,68E-07 | 0,11007 | 576,75 | 64,25 | 29 |
| pavoniella2 | pavoniella5 | pavonia3 | 0,37788 | 3,88439 | 0,000103 | 0,115412 | 685,25 | 74,75 | 33,75 |
| pavonia5 | pavonia1 | pavoniella10 | 0,377588 | 6,13099 | 8,73E-10 | 0,092593 | 208,125 | 141,375 | 63,875 |
| pavoniella13 | pavoniella11 | pavonia4 | 0,377163 | 3,30625 | 0,000946 | 0,081832 | 670,25 | 49,75 | 22,5 |
| pavonia5 | pavonia2 | pavoniella12 | 0,37707 | 5,72133 | 1,06E-08 | 0,083853 | 212,375 | 135,125 | 61,125 |
| pavonia10 | pavonia7 | pavoniella6 | 0,376754 | 6,99823 | 2,59E-12 | 0,126557 | 223,75 | 171,75 | 77,75 |
| pavoniella12 | pavoniella11 | pavonia5 | 0,376569 | 2,5872 | 0,009676 | 0,080286 | 692,625 | 41,125 | 18,625 |
| pavonia5 | pavonia10 | pavoniella6 | 0,375847 | 7,18508 | 6,72E-13 | 0,101617 | 204,875 | 152,375 | 69,125 |
| pavonia5 | pavonia9 | pavoniella8 | 0,375138 | 4,62448 | 3,76E-06 | 0,09446 | 204,5 | 156,25 | 71 |
| pavonia5 | pavonia1 | pavoniella7 | 0,375 | 6,92628 | 4,32E-12 | 0,091973 | 230,25 | 151,25 | 68,75 |
| pavonia5 | pavonia10 | pavoniella1 | 0,374083 | 5,5084 | 3,62E-08 | 0,098869 | 187 | 140,5 | 64 |
| pavoniella13 | pavoniella8 | pavoniella6 | 0,373882 | 5,64976 | 1,61E-08 | 0,324031 | 121,5 | 96 | 43,75 |
| pavonia1 | pavonia7 | pavoniella9 | 0,373232 | 6,83911 | 7,97E-12 | 0,112238 | 220,5 | 157,75 | 72 |
| pavonia5 | pavonia10 | pavoniella3 | 0,373099 | 6,05435 | 1,41E-09 | 0,109284 | 191,5 | 146,75 | 67 |
| pavonia5 | pavonia2 | pavoniella4 | 0,372716 | 7,02184 | 2,19E-12 | 0,113417 | 185,375 | 140,875 | 64,375 |
| pavonia5 | pavonia2 | pavoniella11 | 0,372229 | 5,18399 | 2,17E-07 | 0,098033 | 200,5 | 147 | 67,25 |
| pavoniella12 | pavoniella8 | pavoniella6 | 0,371795 | 5,41619 | 6,09E-08 | 0,320695 | 124,375 | 93,625 | 42,875 |
| pavoniella2 | pavoniella14 | pavonia3 | 0,371728 | 5,98376 | 2,18E-09 | 0,107332 | 613 | 65,5 | 30 |
| pavoniella10 | pavoniella14 | pavonia4 | 0,371274 | 5,60488 | 2,08E-08 | 0,111292 | 560,5 | 63,25 | 29 |
| pavoniella12 | pavoniella4 | pavonia10 | 0,371191 | 4,19702 | 2,70E-05 | 0,095171 | 531,875 | 61,875 | 28,375 |
| pavonia1 | pavonia7 | pavoniella13 | 0,37069 | 6,63136 | 3,33E-11 | 0,101058 | 234,25 | 159 | 73 |
| pavoniella12 | pavoniella3 | pavonia4 | 0,370253 | 4,40913 | 1,04E-05 | 0,090909 | 608,625 | 54,125 | 24,875 |
| pavoniella14 | pavonia7 | pavonia5 | 0,369008 | 5,5626 | 2,66E-08 | 0,345873 | 202 | 153,5 | 70,75 |
| pavonia2 | pavonia7 | pavoniella13 | 0,368138 | 5,53705 | 3,08E-08 | 0,101665 | 222,875 | 158,875 | 73,375 |
| pavonia4 | pavonia7 | pavoniella5 | 0,36803 | 6,95878 | 3,43E-12 | 0,136317 | 186,75 | 184 | 85 |
| pavonia5 | pavonia2 | pavoniella8 | 0,367844 | 6,21902 | 5,00E-10 | 0,083587 | 221,375 | 140,375 | 64,875 |
| pavonia5 | pavonia2 | pavoniella14 | 0,367647 | 6,4919 | 8,48E-11 | 0,111193 | 183 | 139,5 | 64,5 |
| pavonia5 | pavonia3 | pavoniella13 | 0,367424 | 6,50303 | 7,87E-11 | 0,078691 | 217,875 | 135,375 | 62,625 |
| pavonia5 | pavonia10 | pavoniella14 | 0,367206 | 6,57544 | 4,85E-11 | 0,116101 | 181,75 | 148 | 68,5 |
| pavoniella3 | pavonia7 | pavonia6 | 0,367155 | 6,72744 | 1,73E-11 | 0,340116 | 178,875 | 163,375 | 75,625 |
| pavoniella2 | pavoniella14 | pavonia10 | 0,366048 | 4,91222 | 9,01E-07 | 0,099067 | 591,125 | 64,375 | 29,875 |
| pavoniella7 | pavoniella14 | pavonia4 | 0,365196 | 4,27424 | 1,92E-05 | 0,115863 | 585,625 | 69,625 | 32,375 |
| pavoniella13 | pavoniella5 | pavonia1 | 0,364066 | 3,67833 | 0,000235 | 0,102735 | 589,125 | 72,125 | 33,625 |
| pavonia5 | pavonia2 | pavoniella5 | 0,363942 | 6,23404 | 4,55E-10 | 0,11002 | 187,5 | 152,25 | 71 |
| pavonia9 | pavonia7 | pavoniella6 | 0,363918 | 5,34075 | 9,26E-08 | 0,122911 | 235,875 | 165,375 | 77,125 |
| pavonia2 | pavonia7 | pavoniella8 | 0,363823 | 5,8843 | 4,00E-09 | 0,104672 | 229,125 | 165,875 | 77,375 |
| pavonia5 | pavonia8 | pavoniella1 | 0,363636 | 6,39442 | 1,61E-10 | 0,104392 | 166 | 153,75 | 71,75 |
| pavoniella4 | pavoniella11 | pavoniella6 | 0,362456 | 5,34427 | 9,08E-08 | 0,284787 | 147,5 | 144,25 | 67,5 |
| pavoniella14 | pavoniella11 | pavoniella6 | 0,362408 | 5,31057 | 1,09E-07 | 0,286686 | 148,125 | 138,625 | 64,875 |
| pavonia2 | pavonia7 | pavoniella10 | 0,361392 | 5,5564 | 2,75E-08 | 0,107728 | 198,875 | 151,625 | 71,125 |
| pavonia5 | pavonia9 | pavoniella1 | 0,361364 | 4,57364 | 4,79E-06 | 0,104126 | 167,75 | 149,75 | 70,25 |
| pavoniella12 | pavoniella5 | pavonia6 | 0,361111 | 5,65849 | 1,53E-08 | 0,118397 | 584,5 | 61,25 | 28,75 |
| pavoniella9 | pavoniella2 | pavoniella5 | 0,361111 | 6,03247 | 1,61E-09 | 0,395137 | 135 | 122,5 | 57,5 |
| pavonia1 | pavonia7 | pavoniella10 | 0,36071 | 6,96988 | 3,17E-12 | 0,106978 | 219 | 153,25 | 72 |
| pavonia5 | pavonia2 | pavoniella7 | 0,360617 | 5,93567 | 2,93E-09 | 0,086733 | 224,375 | 143,375 | 67,375 |
| pavonia10 | pavonia7 | pavoniella8 | 0,360572 | 6,13483 | 8,52E-10 | 0,104314 | 248,75 | 166,5 | 78,25 |
| pavoniella4 | pavonia7 | pavonia5 | 0,360169 | 4,88052 | 1,06E-06 | 0,344478 | 196,75 | 160,5 | 75,5 |
| pavoniella7 | pavoniella5 | pavonia4 | 0,359813 | 4,39158 | 1,13E-05 | 0,115097 | 642,5 | 72,75 | 34,25 |
| pavoniella5 | pavonia7 | pavonia5 | 0,359743 | 7,06319 | 1,63E-12 | 0,337688 | 217,5 | 158,75 | 74,75 |
| pavoniella9 | pavoniella5 | pavonia4 | 0,359375 | 4,62915 | 3,67E-06 | 0,11147 | 591 | 65,25 | 30,75 |
| pavoniella9 | pavoniella14 | pavonia4 | 0,359043 | 4,28824 | 1,80E-05 | 0,111663 | 556,375 | 63,875 | 30,125 |
| pavoniella6 | pavoniella8 | pavoniella2 | 0,358726 | 5,05806 | 4,24E-07 | 0,276119 | 145,875 | 122,625 | 57,875 |
| pavoniella2 | pavoniella14 | pavonia2 | 0,358601 | 4,30367 | 1,68E-05 | 0,09375 | 588,75 | 58,25 | 27,5 |
| pavoniella2 | pavoniella14 | pavonia1 | 0,358586 | 5,14204 | 2,72E-07 | 0,101792 | 601,75 | 67,25 | 31,75 |
| pavoniella13 | pavoniella4 | pavonia3 | 0,358575 | 4,77389 | 1,81E-06 | 0,117176 | 558,5 | 76,25 | 36 |
| pavoniella7 | pavoniella5 | pavonia2 | 0,358491 | 4,23296 | 2,31E-05 | 0,111355 | 594,75 | 72 | 34 |
| pavonia9 | pavonia7 | pavoniella11 | 0,358289 | 5,78038 | 7,45E-09 | 0,109656 | 240,5 | 158,75 | 75 |
| pavonia3 | pavonia7 | pavoniella11 | 0,35818 | 5,58609 | 2,32E-08 | 0,117386 | 219,875 | 175,375 | 82,875 |
| pavoniella13 | pavoniella4 | pavonia5 | 0,357513 | 3,47633 | 0,000508 | 0,122232 | 616,75 | 65,5 | 31 |
| pavonia1 | pavonia7 | pavoniella7 | 0,357006 | 7,16842 | 7,59E-13 | 0,112285 | 240 | 176,75 | 83,75 |
| pavoniella13 | pavoniella3 | pavonia4 | 0,356287 | 4,46837 | 7,88E-06 | 0,091468 | 619,875 | 56,625 | 26,875 |
| pavoniella7 | pavoniella14 | pavonia3 | 0,355814 | 5,80435 | 6,46E-09 | 0,115559 | 571,375 | 72,875 | 34,625 |
| pavonia5 | pavonia1 | pavoniella3 | 0,355556 | 5,29089 | 1,22E-07 | 0,098833 | 199,5 | 137,25 | 65,25 |
| pavoniella7 | pavoniella11 | pavonia5 | 0,355311 | 3,00899 | 0,002621 | 0,084642 | 724,75 | 46,25 | 22 |
| pavoniella10 | pavoniella14 | pavonia5 | 0,355224 | 4,32039 | 1,56E-05 | 0,118173 | 613,25 | 56,75 | 27 |
| pavonia10 | pavonia7 | pavoniella12 | 0,355191 | 7,9664 | 1,63E-15 | 0,098575 | 241 | 155 | 73,75 |
| pavoniella13 | pavoniella6 | pavonia7 | 0,355014 | 4,13491 | 3,55E-05 | 0,097325 | 569,25 | 62,5 | 29,75 |
| pavonia1 | pavonia7 | pavoniella8 | 0,354935 | 5,67794 | 1,36E-08 | 0,103458 | 242,625 | 169,875 | 80,875 |
| pavoniella8 | pavoniella14 | pavonia2 | 0,354331 | 5,53706 | 3,08E-08 | 0,10235 | 553,25 | 64,5 | 30,75 |
| pavonia10 | pavonia7 | pavoniella11 | 0,354189 | 6,13621 | 8,45E-10 | 0,109508 | 230,625 | 159,625 | 76,125 |
| pavoniella1 | pavonia7 | pavonia6 | 0,354037 | 6,56174 | 5,32E-11 | 0,333333 | 183 | 163,5 | 78 |
| pavonia5 | pavonia1 | pavoniella8 | 0,353286 | 5,55123 | 2,84E-08 | 0,081638 | 231,125 | 144,125 | 68,875 |
| pavoniella13 | pavoniella5 | pavonia6 | 0,352941 | 6,02321 | 1,71E-09 | 0,123104 | 593,625 | 66,125 | 31,625 |
| pavoniella9 | pavoniella5 | pavonia6 | 0,352778 | 4,51124 | 6,44E-06 | 0,120151 | 579,625 | 60,875 | 29,125 |
| pavonia5 | pavonia1 | pavoniella9 | 0,352638 | 4,67582 | 2,93E-06 | 0,082855 | 212,375 | 131,375 | 62,875 |
| pavonia4 | pavonia7 | pavoniella14 | 0,352571 | 6,16727 | 6,95E-10 | 0,129082 | 170,625 | 161,125 | 77,125 |
| pavoniella9 | pavoniella14 | pavonia6 | 0,352239 | 3,70197 | 0,000214 | 0,114341 | 547,625 | 56,625 | 27,125 |
| pavonia3 | pavonia7 | pavoniella2 | 0,35206 | 6,00637 | 1,90E-09 | 0,11339 | 241,5 | 180,5 | 86,5 |
| pavonia5 | pavonia1 | pavoniella12 | 0,35197 | 4,98325 | 6,25E-07 | 0,077765 | 222,75 | 133 | 63,75 |
| pavoniella10 | pavoniella2 | pavoniella14 | 0,351562 | 6,18412 | 6,24E-10 | 0,375626 | 145,125 | 108,125 | 51,875 |
| pavoniella13 | pavoniella2 | pavoniella14 | 0,351477 | 4,44934 | 8,61E-06 | 0,3711 | 136,125 | 108,625 | 52,125 |
| pavonia5 | pavonia8 | pavoniella2 | 0,351269 | 6,45732 | 1,07E-10 | 0,097355 | 191,375 | 166,375 | 79,875 |
| pavonia10 | pavonia7 | pavoniella13 | 0,350972 | 6,80185 | 1,03E-11 | 0,096325 | 237,125 | 156,375 | 75,125 |
| pavonia5 | pavonia8 | pavoniella3 | 0,350664 | 6,72317 | 1,78E-11 | 0,106198 | 171,875 | 152,625 | 73,375 |
| pavonia5 | pavonia8 | pavoniella11 | 0,350526 | 6,31505 | 2,70E-10 | 0,098346 | 179,625 | 160,375 | 77,125 |
| pavonia3 | pavonia7 | pavoniella3 | 0,349896 | 6,0239 | 1,70E-09 | 0,123403 | 202,5 | 163 | 78,5 |
| pavonia5 | pavonia10 | pavoniella4 | 0,349398 | 5,65928 | 1,52E-08 | 0,106578 | 187,25 | 140 | 67,5 |
| pavoniella7 | pavoniella14 | pavonia2 | 0,349246 | 4,43521 | 9,20E-06 | 0,104433 | 549,625 | 67,125 | 32,375 |
| pavonia2 | pavonia7 | pavoniella12 | 0,349241 | 5,24883 | 1,53E-07 | 0,097932 | 221,5 | 155,5 | 75 |
| pavonia5 | pavonia10 | pavoniella12 | 0,349091 | 5,70003 | 1,20E-08 | 0,080831 | 217,375 | 139,125 | 67,125 |
| pavoniella7 | pavoniella14 | pavonia5 | 0,348774 | 3,80328 | 0,000143 | 0,115628 | 634,125 | 61,875 | 29,875 |
| pavoniella10 | pavoniella5 | pavonia3 | 0,348519 | 4,36142 | 1,29E-05 | 0,118881 | 584,75 | 74 | 35,75 |
| pavoniella13 | pavoniella6 | pavonia10 | 0,348485 | 2,60244 | 0,009256 | 0,064561 | 638 | 44,5 | 21,5 |
| pavoniella10 | pavoniella2 | pavoniella3 | 0,34816 | 6,29859 | 3,00E-10 | 0,403915 | 120,875 | 109,875 | 53,125 |
| pavoniella9 | pavoniella5 | pavonia3 | 0,347722 | 3,79987 | 0,000145 | 0,111196 | 575 | 70,25 | 34 |
| pavonia5 | pavonia3 | pavoniella9 | 0,347368 | 5,03194 | 4,86E-07 | 0,079375 | 203,5 | 128 | 62 |
| pavonia5 | pavonia1 | pavoniella2 | 0,347144 | 4,85385 | 1,21E-06 | 0,088344 | 220,875 | 150,375 | 72,875 |
| pavoniella13 | pavoniella4 | pavonia4 | 0,346835 | 3,90567 | 9,40E-05 | 0,105955 | 582,5 | 66,5 | 32,25 |
| pavoniella12 | pavoniella1 | pavonia2 | 0,34657 | 3,83718 | 0,000124 | 0,076677 | 600,125 | 46,625 | 22,625 |
| pavonia5 | pavonia1 | pavoniella1 | 0,346299 | 4,46664 | 7,95E-06 | 0,090226 | 196,875 | 134,125 | 65,125 |
| pavoniella10 | pavoniella14 | pavonia3 | 0,346062 | 5,17708 | 2,25E-07 | 0,117886 | 547,75 | 70,5 | 34,25 |
| pavoniella12 | pavoniella5 | pavonia10 | 0,345455 | 3,91415 | 9,07E-05 | 0,092042 | 571,75 | 64,75 | 31,5 |
| pavonia9 | pavonia7 | pavoniella13 | 0,344676 | 5,20593 | 1,93E-07 | 0,094635 | 250,375 | 153,125 | 74,625 |
| pavoniella6 | pavoniella1 | pavoniella4 | 0,344527 | 4,95206 | 7,34E-07 | 0,422901 | 138,125 | 135,125 | 65,875 |
| pavoniella13 | pavoniella4 | pavonia10 | 0,344173 | 3,68661 | 0,000227 | 0,090974 | 536 | 62 | 30,25 |
| pavoniella8 | pavoniella11 | pavonia5 | 0,344 | 2,9065 | 0,003655 | 0,074848 | 734,75 | 42 | 20,5 |
| pavonia10 | pavonia7 | pavoniella10 | 0,343956 | 6,22051 | 4,96E-10 | 0,104368 | 217,875 | 152,875 | 74,625 |
| pavonia5 | pavonia1 | pavoniella11 | 0,34386 | 4,61626 | 3,91E-06 | 0,089253 | 208,125 | 143,625 | 70,125 |
| pavoniella12 | pavoniella4 | pavonia5 | 0,343832 | 3,36266 | 0,000772 | 0,119854 | 612,25 | 64 | 31,25 |
| pavonia1 | pavonia7 | pavoniella1 | 0,34375 | 5,33087 | 9,77E-08 | 0,109843 | 216 | 150,5 | 73,5 |
| pavonia10 | pavonia7 | pavoniella9 | 0,343578 | 5,90653 | 3,49E-09 | 0,104473 | 226 | 153 | 74,75 |
| pavonia1 | pavonia7 | pavoniella12 | 0,343042 | 6,5549 | 5,57E-11 | 0,095352 | 235,875 | 155,625 | 76,125 |
| pavonia5 | pavonia10 | pavoniella8 | 0,342625 | 5,71469 | 1,10E-08 | 0,081447 | 223,5 | 144,5 | 70,75 |
| pavonia4 | pavonia7 | pavoniella4 | 0,341141 | 9,51516 | 2,30E-16 | 0,125798 | 188,125 | 164,625 | 80,875 |
| pavoniella12 | pavoniella2 | pavoniella5 | 0,340366 | 4,27161 | 1,94E-05 | 0,367223 | 132,625 | 119,125 | 58,625 |
| pavonia5 | pavonia2 | pavoniella2 | 0,340254 | 4,9878 | 6,11E-07 | 0,086714 | 215,75 | 145,25 | 71,5 |
| pavonia5 | pavonia3 | pavoniella1 | 0,34018 | 4,98755 | 6,11E-07 | 0,084584 | 189,25 | 130,5 | 64,25 |
| pavoniella12 | pavoniella1 | pavonia4 | 0,34 | 2,97064 | 0,002972 | 0,068493 | 637,375 | 41,875 | 20,625 |
| pavoniella12 | pavoniella2 | pavoniella14 | 0,339344 | 4,89597 | 9,78E-07 | 0,352041 | 141,375 | 102,125 | 50,375 |
| pavoniella8 | pavoniella4 | pavonia10 | 0,339332 | 4,07568 | 4,59E-05 | 0,09173 | 560,875 | 65,125 | 32,125 |
| pavoniella9 | pavoniella2 | pavoniella14 | 0,338141 | 4,58038 | 4,64E-06 | 0,358234 | 138,625 | 104,375 | 51,625 |
| pavoniella10 | pavoniella14 | pavonia1 | 0,338061 | 4,25969 | 2,05E-05 | 0,110254 | 532 | 70,75 | 35 |
| pavonia8 | pavonia7 | pavoniella10 | 0,337808 | 5,53944 | 3,03E-08 | 0,098757 | 246,5 | 149,5 | 74 |
| pavoniella8 | pavoniella4 | pavonia2 | 0,337408 | 4,23549 | 2,28E-05 | 0,103448 | 560,625 | 68,375 | 33,875 |
| pavonia9 | pavonia7 | pavoniella8 | 0,337185 | 4,90087 | 9,54E-07 | 0,095935 | 257,625 | 159,125 | 78,875 |
| pavoniella9 | pavoniella14 | pavonia10 | 0,336538 | 4,35091 | 1,36E-05 | 0,106141 | 523,75 | 69,5 | 34,5 |
| pavoniella2 | pavoniella4 | pavonia2 | 0,335917 | 4,62806 | 3,69E-06 | 0,097744 | 601,375 | 64,625 | 32,125 |
| pavonia2 | pavonia7 | pavoniella11 | 0,335498 | 5,70203 | 1,18E-08 | 0,102785 | 210 | 154,25 | 76,75 |
| pavonia5 | pavonia3 | pavoniella6 | 0,335351 | 5,69265 | 1,25E-08 | 0,08399 | 206,375 | 137,875 | 68,625 |
| pavoniella14 | pavoniella1 | pavoniella6 | 0,33462 | 5,15921 | 2,48E-07 | 0,265577 | 130,875 | 129,625 | 64,625 |
| pavoniella8 | pavoniella11 | pavonia4 | 0,334495 | 2,91986 | 0,003502 | 0,071376 | 702,875 | 47,875 | 23,875 |
| pavoniella12 | pavoniella3 | pavonia10 | 0,334375 | 4,26916 | 1,96E-05 | 0,075887 | 562,125 | 53,375 | 26,625 |
| pavonia9 | pavonia7 | pavoniella7 | 0,333667 | 5,07479 | 3,88E-07 | 0,103384 | 254,375 | 166,375 | 83,125 |
| pavonia5 | pavonia1 | pavoniella5 | 0,332953 | 5,47248 | 4,44E-08 | 0,097463 | 192,125 | 146,125 | 73,125 |
| pavoniella6 | pavoniella8 | pavoniella14 | 0,332734 | 3,73198 | 0,00019 | 0,254821 | 202,375 | 92,625 | 46,375 |
| pavonia9 | pavonia7 | pavoniella2 | 0,332636 | 5,07587 | 3,86E-07 | 0,100284 | 258,25 | 159,25 | 79,75 |
| pavoniella8 | pavoniella4 | pavonia3 | 0,332599 | 4,60984 | 4,03E-06 | 0,109978 | 585,125 | 75,625 | 37,875 |
| pavoniella5 | pavoniella2 | pavoniella6 | 0,332126 | 4,75509 | 1,98E-06 | 0,25369 | 167,625 | 137,875 | 69,125 |
| pavoniella2 | pavoniella14 | pavonia4 | 0,331378 | 4,27221 | 1,94E-05 | 0,08954 | 626,75 | 56,75 | 28,5 |
| pavonia9 | pavonia7 | pavoniella12 | 0,331133 | 4,90896 | 9,16E-07 | 0,092077 | 245,5 | 151,25 | 76 |
| pavonia5 | pavonia10 | pavoniella7 | 0,33 | 5,55183 | 2,83E-08 | 0,083474 | 223,875 | 149,625 | 75,375 |
| pavoniella6 | pavoniella8 | pavoniella3 | 0,329932 | 5,20743 | 1,91E-07 | 0,257979 | 192,75 | 97,75 | 49,25 |
| pavoniella12 | pavoniella6 | pavonia10 | 0,329749 | 3,15097 | 0,001627 | 0,064471 | 620,875 | 46,375 | 23,375 |
| pavonia5 | pavonia10 | pavoniella11 | 0,329558 | 4,78893 | 1,68E-06 | 0,087703 | 204,75 | 146,75 | 74 |
| pavoniella7 | pavoniella5 | pavonia3 | 0,329032 | 3,87973 | 0,000105 | 0,110469 | 621,75 | 77,25 | 39 |
| pavoniella9 | pavoniella5 | pavonia1 | 0,328467 | 3,70878 | 0,000208 | 0,095004 | 563,5 | 68,25 | 34,5 |
| pavonia5 | pavonia3 | pavoniella12 | 0,328105 | 5,03193 | 4,86E-07 | 0,070367 | 212 | 127 | 64,25 |
| pavonia9 | pavonia7 | pavoniella10 | 0,327684 | 5,15171 | 2,58E-07 | 0,098305 | 225,375 | 146,875 | 74,375 |
| pavoniella9 | pavoniella5 | pavonia2 | 0,327543 | 3,91111 | 9,19E-05 | 0,105347 | 553,625 | 66,875 | 33,875 |
| pavonia5 | pavonia3 | pavoniella3 | 0,327411 | 5,03435 | 4,79E-07 | 0,087815 | 188,75 | 130,75 | 66,25 |
| pavoniella8 | pavoniella5 | pavonia1 | 0,326241 | 3,56696 | 0,000361 | 0,090551 | 624,875 | 70,125 | 35,625 |
| pavonia1 | pavonia7 | pavoniella2 | 0,325674 | 5,79719 | 6,74E-09 | 0,099939 | 239,625 | 165,875 | 84,375 |
| pavonia5 | pavonia3 | pavoniella10 | 0,32567 | 5,01723 | 5,24E-07 | 0,076074 | 201,25 | 129,75 | 66 |
| pavoniella13 | pavoniella2 | pavoniella5 | 0,325611 | 4,73367 | 2,20E-06 | 0,354839 | 139,5 | 128,75 | 65,5 |
| pavonia5 | pavonia9 | pavoniella2 | 0,325557 | 3,70454 | 0,000212 | 0,090427 | 198,5 | 156,25 | 79,5 |
| pavonia5 | pavonia9 | pavoniella4 | 0,325556 | 4,08633 | 4,38E-05 | 0,109003 | 170,875 | 149,125 | 75,875 |
| pavoniella7 | pavoniella5 | pavonia6 | 0,325459 | 3,7664 | 0,000166 | 0,113866 | 607,125 | 63,125 | 32,125 |
| pavonia5 | pavonia3 | pavoniella14 | 0,325213 | 5,33119 | 9,76E-08 | 0,096739 | 176,25 | 136 | 69,25 |
| pavonia2 | pavonia7 | pavoniella2 | 0,324974 | 5,54556 | 2,93E-08 | 0,098449 | 229 | 158,5 | 80,75 |
| pavonia5 | pavonia8 | pavoniella5 | 0,32495 | 6,15073 | 7,71E-10 | 0,105246 | 169,125 | 164,625 | 83,875 |
| pavonia3 | pavonia7 | pavoniella1 | 0,324847 | 4,46734 | 7,92E-06 | 0,111034 | 205,875 | 162,625 | 82,875 |
| pavoniella12 | pavoniella6 | pavonia2 | 0,324818 | 2,33485 | 0,019551 | 0,067069 | 612,375 | 45,375 | 23,125 |
| pavoniella6 | pavoniella8 | pavoniella11 | 0,324575 | 4,27336 | 1,93E-05 | 0,231278 | 145,875 | 107,125 | 54,625 |
| pavonia1 | pavonia7 | pavoniella11 | 0,324435 | 5,33274 | 9,67E-08 | 0,102631 | 225,75 | 161,25 | 82,25 |
| pavonia5 | pavonia8 | pavoniella14 | 0,324239 | 6,15377 | 7,57E-10 | 0,109847 | 167,25 | 157,75 | 80,5 |
| pavoniella4 | pavonia7 | pavonia6 | 0,324074 | 6,65017 | 2,93E-11 | 0,3125 | 171,625 | 160,875 | 82,125 |
| pavoniella13 | pavoniella6 | pavonia2 | 0,32342 | 2,15387 | 0,031251 | 0,064444 | 633,5 | 44,5 | 22,75 |
| pavoniella3 | pavoniella2 | pavoniella6 | 0,32341 | 4,72156 | 2,34E-06 | 0,2364 | 160,75 | 122,25 | 62,5 |
| pavoniella7 | pavoniella2 | pavoniella14 | 0,323144 | 4,97308 | 6,59E-07 | 0,360976 | 154,625 | 113,625 | 58,125 |
| pavoniella13 | pavoniella3 | pavonia2 | 0,322946 | 4,1192 | 3,80E-05 | 0,085586 | 574,875 | 58,375 | 29,875 |
| pavoniella12 | pavoniella2 | pavonia4 | 0,32287 | 3,08458 | 0,002038 | 0,053137 | 692,625 | 36,875 | 18,875 |
| pavoniella6 | pavoniella9 | pavoniella2 | 0,322422 | 3,99454 | 6,48E-05 | 0,238499 | 124,5 | 101 | 51,75 |
| pavonia5 | pavonia3 | pavoniella4 | 0,322222 | 4,67909 | 2,88E-06 | 0,094875 | 184,125 | 133,875 | 68,625 |
| pavoniella10 | pavoniella2 | pavoniella5 | 0,321023 | 5,07444 | 3,89E-07 | 0,347692 | 137,5 | 116,25 | 59,75 |
| pavoniella12 | pavoniella4 | pavonia3 | 0,320843 | 3,85843 | 0,000114 | 0,103553 | 552,75 | 70,5 | 36,25 |
| pavonia10 | pavonia7 | pavoniella2 | 0,320419 | 6,69859 | 2,10E-11 | 0,096866 | 244,875 | 157,625 | 81,125 |
| pavoniella12 | pavoniella3 | pavonia2 | 0,320359 | 3,77089 | 0,000163 | 0,082371 | 565,875 | 55,125 | 28,375 |
| pavonia9 | pavonia7 | pavoniella1 | 0,320356 | 4,62068 | 3,82E-06 | 0,104499 | 225,625 | 148,375 | 76,375 |
| pavoniella7 | pavoniella14 | pavonia1 | 0,320175 | 4,31968 | 1,56E-05 | 0,104735 | 553 | 75,25 | 38,75 |
| pavonia5 | pavonia1 | pavoniella4 | 0,32 | 4,56698 | 4,95E-06 | 0,094152 | 191,5 | 132 | 68 |
| pavonia1 | pavonia7 | pavoniella3 | 0,32 | 5,261 | 1,43E-07 | 0,110037 | 213,125 | 152,625 | 78,625 |
| pavoniella2 | pavoniella4 | pavonia3 | 0,32 | 3,83411 | 0,000126 | 0,098337 | 627,125 | 70,125 | 36,125 |
| pavoniella7 | pavoniella2 | pavoniella5 | 0,319703 | 5,45995 | 4,76E-08 | 0,353425 | 159,375 | 133,125 | 68,625 |
| pavoniella8 | pavoniella5 | pavonia6 | 0,319149 | 4,06072 | 4,89E-05 | 0,107239 | 616,25 | 62 | 32 |
| pavoniella6 | pavoniella8 | pavoniella1 | 0,318865 | 5,15379 | 2,55E-07 | 0,265278 | 138,75 | 98,75 | 51 |
| pavoniella7 | pavoniella3 | pavonia2 | 0,318841 | 4,53525 | 5,75E-06 | 0,082956 | 589,125 | 56,875 | 29,375 |
| pavonia5 | pavonia3 | pavoniella5 | 0,318644 | 5,24952 | 1,52E-07 | 0,093969 | 184,375 | 145,875 | 75,375 |
| pavoniella13 | pavoniella2 | pavoniella3 | 0,318608 | 5,15765 | 2,50E-07 | 0,389525 | 129,125 | 123,125 | 63,625 |
| pavoniella13 | pavoniella1 | pavoniella4 | 0,318182 | 5,35756 | 8,44E-08 | 0,376083 | 135,625 | 112,375 | 58,125 |
| pavonia5 | pavonia9 | pavoniella11 | 0,317359 | 4,17305 | 3,01E-05 | 0,091159 | 182,375 | 154,625 | 80,125 |
| pavoniella7 | pavoniella11 | pavonia4 | 0,317308 | 3,35742 | 0,000787 | 0,073771 | 693,375 | 51,375 | 26,625 |
| pavoniella13 | pavoniella6 | pavonia9 | 0,316901 | 2,57018 | 0,010165 | 0,058027 | 623,25 | 46,75 | 24,25 |
| pavoniella13 | pavoniella6 | pavonia3 | 0,316726 | 2,25124 | 0,02437 | 0,063526 | 658 | 46,25 | 24 |
| pavoniella10 | pavoniella8 | pavoniella6 | 0,316375 | 5,00367 | 5,62E-07 | 0,326765 | 115,75 | 103,5 | 53,75 |
| pavoniella13 | pavoniella3 | pavonia10 | 0,315634 | 3,72125 | 0,000198 | 0,076813 | 571 | 55,75 | 29 |
| pavoniella9 | pavoniella14 | pavonia2 | 0,315508 | 3,51952 | 0,000432 | 0,09696 | 517,75 | 61,5 | 32 |
| pavoniella6 | pavoniella8 | pavoniella5 | 0,315301 | 3,64419 | 0,000268 | 0,24878 | 209,375 | 106,375 | 55,375 |
| pavonia5 | pavonia10 | pavoniella2 | 0,315265 | 4,47791 | 7,54E-06 | 0,083382 | 216,125 | 148,625 | 77,375 |
| pavonia5 | pavonia8 | pavoniella4 | 0,315054 | 7,08559 | 1,38E-12 | 0,105358 | 165,625 | 152,875 | 79,625 |
| pavoniella2 | pavoniella5 | pavonia6 | 0,314286 | 3,07857 | 0,00208 | 0,099099 | 657,25 | 57,5 | 30 |
| pavonia5 | pavonia3 | pavoniella7 | 0,31295 | 5,14506 | 2,67E-07 | 0,073438 | 218,625 | 136,875 | 71,625 |
| pavoniella13 | pavoniella2 | pavoniella4 | 0,312775 | 4,04985 | 5,13E-05 | 0,337559 | 152,5 | 111,75 | 58,5 |
| pavoniella12 | pavoniella9 | pavoniella6 | 0,312721 | 3,67141 | 0,000241 | 0,302048 | 104,875 | 92,875 | 48,625 |
| pavonia8 | pavonia7 | pavoniella7 | 0,311891 | 5,73441 | 9,79E-09 | 0,096415 | 267,75 | 168,25 | 88,25 |
| pavoniella14 | pavonia7 | pavonia6 | 0,311688 | 5,62021 | 1,91E-08 | 0,30094 | 173,75 | 151,5 | 79,5 |
| pavoniella13 | pavoniella11 | pavoniella4 | 0,311653 | 5,11845 | 3,08E-07 | 0,383333 | 138,5 | 121 | 63,5 |
| pavoniella10 | pavoniella11 | pavoniella5 | 0,310576 | 5,16455 | 2,41E-07 | 0,370607 | 122,625 | 122,375 | 64,375 |
| pavoniella12 | pavoniella14 | pavonia6 | 0,310541 | 4,89756 | 9,70E-07 | 0,10322 | 552,5 | 57,5 | 30,25 |
| pavonia9 | pavonia7 | pavoniella9 | 0,309843 | 4,37906 | 1,19E-05 | 0,095124 | 231,625 | 146,375 | 77,125 |
| pavonia8 | pavonia7 | pavoniella6 | 0,309659 | 6,05148 | 1,44E-09 | 0,10835 | 250,625 | 172,875 | 91,125 |
| pavoniella9 | pavoniella11 | pavonia5 | 0,309237 | 2,36734 | 0,017916 | 0,071362 | 676 | 40,75 | 21,5 |
| pavonia8 | pavonia7 | pavoniella3 | 0,308696 | 5,26362 | 1,41E-07 | 0,103273 | 245,25 | 150,5 | 79,5 |
| pavoniella12 | pavoniella2 | pavoniella4 | 0,308682 | 3,22686 | 0,001252 | 0,323232 | 158 | 101,75 | 53,75 |
| pavoniella12 | pavoniella1 | pavonia10 | 0,308642 | 3,01529 | 0,002567 | 0,055556 | 593,25 | 39,75 | 21 |
| pavoniella2 | pavoniella5 | pavonia5 | 0,308383 | 2,85545 | 0,004298 | 0,091882 | 737,875 | 54,625 | 28,875 |
| pavonia5 | pavonia3 | pavoniella8 | 0,308261 | 4,80045 | 1,58E-06 | 0,068568 | 220,125 | 132,625 | 70,125 |
| pavonia9 | pavonia7 | pavoniella14 | 0,308046 | 4,35569 | 1,33E-05 | 0,108283 | 221,5 | 142,25 | 75,25 |
| pavonia5 | pavonia9 | pavoniella5 | 0,308012 | 3,97047 | 7,17E-05 | 0,099899 | 170,375 | 157,125 | 83,125 |
| pavoniella2 | pavoniella5 | pavonia4 | 0,307487 | 3,41979 | 0,000627 | 0,086272 | 712,375 | 61,125 | 32,375 |
| pavonia2 | pavonia7 | pavoniella3 | 0,307263 | 4,99724 | 5,82E-07 | 0,104048 | 198 | 146,25 | 77,5 |
| pavonia8 | pavonia7 | pavoniella13 | 0,307045 | 4,78548 | 1,71E-06 | 0,085631 | 265,125 | 155,375 | 82,375 |
| pavonia5 | pavonia3 | pavoniella11 | 0,306792 | 4,46807 | 7,89E-06 | 0,077976 | 201,5 | 139,5 | 74 |
| pavonia5 | pavonia10 | pavoniella5 | 0,305335 | 4,46614 | 7,96E-06 | 0,090725 | 189,75 | 143,75 | 76,5 |
| pavonia8 | pavonia7 | pavoniella1 | 0,30473 | 4,62622 | 3,72E-06 | 0,096247 | 247,5 | 148,25 | 79 |
| pavoniella7 | pavoniella4 | pavonia2 | 0,304348 | 3,92102 | 8,82E-05 | 0,094382 | 551,25 | 67,5 | 36 |
| pavonia2 | pavonia7 | pavoniella1 | 0,3043 | 4,67059 | 3,00E-06 | 0,099067 | 198,125 | 147,875 | 78,875 |
| pavoniella8 | pavoniella9 | pavonia9 | 0,303797 | 2,61291 | 0,008977 | 0,032877 | 669,25 | 25,75 | 13,75 |
| pavoniella2 | pavoniella4 | pavonia10 | 0,303279 | 3,71093 | 0,000207 | 0,077298 | 598,875 | 59,625 | 31,875 |
| pavonia10 | pavonia7 | pavoniella5 | 0,302743 | 5,12742 | 2,94E-07 | 0,104859 | 219,125 | 154,375 | 82,625 |
| pavoniella11 | pavoniella14 | pavonia10 | 0,302632 | 3,52345 | 0,000426 | 0,085248 | 580,625 | 61,875 | 33,125 |
| pavonia8 | pavonia7 | pavoniella11 | 0,302301 | 5,02381 | 5,07E-07 | 0,092068 | 261,875 | 155,625 | 83,375 |
| pavonia9 | pavonia7 | pavoniella3 | 0,301952 | 4,60946 | 4,04E-06 | 0,100574 | 226,25 | 141,75 | 76 |
| pavoniella12 | pavoniella10 | pavonia10 | 0,301887 | 2,66792 | 0,007632 | 0,03532 | 645,375 | 25,875 | 13,875 |
| pavoniella12 | pavoniella2 | pavoniella3 | 0,30131 | 4,19079 | 2,78E-05 | 0,372302 | 122,25 | 111,75 | 60 |
| pavoniella9 | pavoniella4 | pavonia5 | 0,301047 | 2,9104 | 0,00361 | 0,108696 | 608,875 | 62,125 | 33,375 |
| pavoniella6 | pavoniella7 | pavoniella2 | 0,300671 | 5,60601 | 2,07E-08 | 0,248337 | 136,125 | 121,125 | 65,125 |
| pavonia10 | pavonia7 | pavoniella1 | 0,300443 | 5,48376 | 4,16E-08 | 0,096579 | 216,125 | 146,625 | 78,875 |
| pavoniella13 | pavoniella14 | pavonia6 | 0,3 | 4,33155 | 1,48E-05 | 0,09887 | 557,875 | 56,875 | 30,625 |
| pavoniella9 | pavoniella2 | pavoniella3 | 0,29985 | 4,20153 | 2,65E-05 | 0,359712 | 115,375 | 108,375 | 58,375 |
| pavoniella8 | pavoniella3 | pavonia4 | 0,29878 | 3,16067 | 0,001574 | 0,075501 | 633,5 | 53,25 | 28,75 |
| pavoniella13 | pavoniella11 | pavoniella14 | 0,297753 | 3,82044 | 0,000133 | 0,358108 | 132,25 | 115,5 | 62,5 |
| pavonia8 | pavonia7 | pavoniella12 | 0,297645 | 4,62291 | 3,78E-06 | 0,082886 | 262 | 151,5 | 82 |
| pavoniella13 | pavoniella3 | pavonia3 | 0,297587 | 3,82557 | 0,00013 | 0,081738 | 598,25 | 60,5 | 32,75 |
| pavoniella12 | pavoniella6 | pavonia8 | 0,297578 | 2,43449 | 0,014913 | 0,059889 | 620,125 | 46,875 | 25,375 |
| pavonia1 | pavonia7 | pavoniella5 | 0,297521 | 4,89235 | 9,96E-07 | 0,103859 | 212 | 157 | 85 |
| pavoniella8 | pavoniella6 | pavonia8 | 0,297125 | 2,71681 | 0,006591 | 0,061225 | 710,25 | 50,75 | 27,5 |
| pavonia8 | pavonia7 | pavoniella8 | 0,297087 | 4,70074 | 2,59E-06 | 0,087529 | 278,5 | 167 | 90,5 |
| pavoniella1 | pavoniella14 | pavonia1 | 0,296651 | 4,49416 | 6,98E-06 | 0,094297 | 557,25 | 67,75 | 36,75 |
| pavoniella8 | pavoniella3 | pavonia2 | 0,296512 | 3,53433 | 0,000409 | 0,076865 | 588,75 | 55,75 | 30,25 |
| pavoniella8 | pavoniella4 | pavonia5 | 0,29602 | 3,00881 | 0,002623 | 0,105217 | 645,375 | 65,125 | 35,375 |
| pavoniella8 | pavoniella6 | pavonia7 | 0,295213 | 3,46685 | 0,000527 | 0,081081 | 634,875 | 60,875 | 33,125 |
| pavoniella13 | pavoniella1 | pavonia4 | 0,295203 | 2,66906 | 0,007606 | 0,063847 | 641,625 | 43,875 | 23,875 |
| pavoniella8 | pavoniella6 | pavonia4 | 0,295203 | 2,54377 | 0,010967 | 0,059172 | 753,375 | 43,875 | 23,875 |
| pavoniella6 | pavoniella9 | pavoniella3 | 0,295203 | 3,58181 | 0,000341 | 0,22409 | 171 | 87,75 | 47,75 |
| pavoniella8 | pavoniella4 | pavonia4 | 0,294686 | 3,95617 | 7,62E-05 | 0,093558 | 610,75 | 67 | 36,5 |
| pavoniella13 | pavoniella3 | pavonia5 | 0,294671 | 3,35906 | 0,000782 | 0,084229 | 653,375 | 51,625 | 28,125 |
| pavoniella13 | pavoniella1 | pavonia5 | 0,294355 | 2,73467 | 0,006244 | 0,066123 | 682,375 | 40,125 | 21,875 |
| pavonia5 | pavonia9 | pavoniella14 | 0,294311 | 3,9171 | 8,96E-05 | 0,098391 | 167,875 | 147,875 | 80,625 |
| pavoniella8 | pavoniella14 | pavonia6 | 0,293785 | 3,85259 | 0,000117 | 0,097106 | 567,5 | 57,25 | 31,25 |
| pavonia2 | pavonia7 | pavoniella14 | 0,293772 | 4,11707 | 3,84E-05 | 0,102669 | 184,625 | 137,625 | 75,125 |
| pavoniella12 | pavoniella6 | pavonia7 | 0,292876 | 3,43951 | 0,000583 | 0,08371 | 559 | 61,25 | 33,5 |
| pavoniella9 | pavoniella6 | pavonia4 | 0,292373 | 2,12012 | 0,033996 | 0,055422 | 682,875 | 38,125 | 20,875 |
| pavoniella13 | pavoniella7 | pavoniella6 | 0,292254 | 4,01939 | 5,83E-05 | 0,254992 | 120,5 | 91,75 | 50,25 |
| pavoniella12 | pavoniella11 | pavoniella4 | 0,291971 | 4,39875 | 1,09E-05 | 0,352734 | 144,125 | 110,625 | 60,625 |
| pavonia9 | pavonia7 | pavoniella5 | 0,291005 | 4,73298 | 2,21E-06 | 0,10077 | 232,75 | 152,5 | 83,75 |
| pavoniella10 | pavoniella14 | pavonia10 | 0,29078 | 3,87482 | 0,000107 | 0,095645 | 527,75 | 68,25 | 37,5 |
| pavoniella10 | pavoniella5 | pavonia1 | 0,290557 | 3,10584 | 0,001897 | 0,0874 | 567,375 | 66,625 | 36,625 |
| pavoniella2 | pavoniella14 | pavonia6 | 0,29 | 2,78255 | 0,005393 | 0,083015 | 607,875 | 48,375 | 26,625 |
| pavonia8 | pavonia7 | pavoniella9 | 0,28973 | 4,63822 | 3,51E-06 | 0,088537 | 244,375 | 149,125 | 82,125 |
| pavoniella8 | pavoniella6 | pavonia2 | 0,288809 | 2,2252 | 0,026068 | 0,05878 | 692,875 | 44,625 | 24,625 |
| pavonia10 | pavonia7 | pavoniella3 | 0,288717 | 5,18538 | 2,16E-07 | 0,099051 | 215,375 | 145,625 | 80,375 |
| pavoniella13 | pavoniella6 | pavonia1 | 0,288525 | 2,48677 | 0,012891 | 0,059419 | 656,125 | 49,125 | 27,125 |
| pavoniella13 | pavoniella12 | pavoniella11 | 0,288462 | 2,264 | 0,023574 | 0,079576 | 228,5 | 33,5 | 18,5 |
| pavoniella8 | pavoniella2 | pavoniella14 | 0,288344 | 4,16074 | 3,17E-05 | 0,326389 | 153,25 | 105 | 58 |
| pavoniella12 | pavoniella11 | pavoniella14 | 0,287856 | 3,64278 | 0,00027 | 0,345946 | 138,125 | 107,375 | 59,375 |
| pavoniella9 | pavoniella11 | pavonia4 | 0,287823 | 2,53206 | 0,011339 | 0,063005 | 627,875 | 43,625 | 24,125 |
| pavoniella8 | pavoniella6 | pavonia10 | 0,287719 | 3,04464 | 0,00233 | 0,056396 | 703,125 | 45,875 | 25,375 |
| pavoniella13 | pavoniella5 | pavonia10 | 0,287105 | 3,44785 | 0,000565 | 0,08223 | 583,875 | 66,125 | 36,625 |
| pavonia8 | pavonia7 | pavoniella2 | 0,286437 | 5,09559 | 3,48E-07 | 0,08519 | 275,375 | 158,875 | 88,125 |
| pavoniella10 | pavoniella9 | pavoniella6 | 0,286416 | 3,57529 | 0,00035 | 0,303819 | 100 | 98,25 | 54,5 |
| pavonia5 | pavonia3 | pavoniella2 | 0,286207 | 4,18231 | 2,89E-05 | 0,0709 | 216,375 | 139,875 | 77,625 |
| pavoniella10 | pavoniella11 | pavoniella4 | 0,285714 | 5,43377 | 5,52E-08 | 0,3762 | 142,5 | 110,25 | 61,25 |
| pavoniella5 | pavoniella11 | pavoniella6 | 0,285714 | 5,35755 | 8,44E-08 | 0,220488 | 161,625 | 127,125 | 70,625 |
| pavoniella7 | pavoniella4 | pavonia5 | 0,285377 | 3,5448 | 0,000393 | 0,109107 | 638,125 | 68,125 | 37,875 |
| pavoniella5 | pavonia7 | pavonia6 | 0,284687 | 6,64843 | 2,96E-11 | 0,284979 | 182 | 156,25 | 87 |
| pavoniella6 | pavoniella9 | pavoniella4 | 0,284333 | 4,53719 | 5,70E-06 | 0,219076 | 196,5 | 83 | 46,25 |
| pavoniella7 | pavoniella6 | pavonia4 | 0,284173 | 2,78152 | 0,005411 | 0,05826 | 738,875 | 44,625 | 24,875 |
| pavoniella10 | pavoniella5 | pavonia6 | 0,283784 | 3,56922 | 0,000358 | 0,101942 | 580,875 | 59,375 | 33,125 |
| pavoniella2 | pavoniella6 | pavonia7 | 0,282979 | 3,56363 | 0,000366 | 0,094326 | 570,625 | 75,375 | 42,125 |
| pavoniella7 | pavoniella4 | pavonia10 | 0,282927 | 3,20278 | 0,001361 | 0,081176 | 556,75 | 65,75 | 36,75 |
| pavoniella13 | pavoniella8 | pavoniella11 | 0,28246 | 3,20933 | 0,00133 | 0,156171 | 136,125 | 70,375 | 39,375 |
| pavoniella12 | pavoniella1 | pavoniella4 | 0,282012 | 4,06659 | 4,77E-05 | 0,338828 | 141,125 | 105,125 | 58,875 |
| pavoniella13 | pavoniella9 | pavoniella6 | 0,282007 | 3,19704 | 0,001388 | 0,27395 | 96,625 | 92,625 | 51,875 |
| pavoniella7 | pavoniella6 | pavonia10 | 0,281967 | 3,08333 | 0,002047 | 0,058824 | 691,375 | 48,875 | 27,375 |
| pavoniella10 | pavoniella11 | pavoniella14 | 0,28169 | 3,89922 | 9,65E-05 | 0,34965 | 138,75 | 113,75 | 63,75 |
| pavoniella12 | pavoniella3 | pavonia5 | 0,281457 | 3,07742 | 0,002088 | 0,078269 | 644,625 | 48,375 | 27,125 |
| pavoniella6 | pavoniella7 | pavoniella3 | 0,281399 | 4,83654 | 1,32E-06 | 0,237265 | 184,25 | 100,75 | 56,5 |
| pavoniella12 | pavoniella1 | pavonia5 | 0,281385 | 2,68963 | 0,007153 | 0,061205 | 675,25 | 37 | 20,75 |
| pavoniella7 | pavoniella2 | pavoniella3 | 0,280585 | 4,41846 | 9,94E-06 | 0,362543 | 135,875 | 120,375 | 67,625 |
| pavonia2 | pavonia7 | pavoniella5 | 0,279958 | 5,19251 | 2,07E-07 | 0,098951 | 196,875 | 150,875 | 84,875 |
| pavonia6 | pavonia8 | pavoniella9 | 0,279863 | 4,01169 | 6,03E-05 | 0,07938 | 172,375 | 140,625 | 79,125 |
| pavoniella10 | pavoniella11 | pavonia5 | 0,279693 | 2,17989 | 0,029266 | 0,070463 | 694,75 | 41,75 | 23,5 |
| pavoniella8 | pavoniella5 | pavonia10 | 0,279621 | 3,63322 | 0,00028 | 0,079838 | 618 | 67,5 | 38 |
| pavoniella7 | pavoniella4 | pavonia3 | 0,278761 | 3,75252 | 0,000175 | 0,092852 | 576,25 | 72,25 | 40,75 |
| pavoniella7 | pavoniella14 | pavonia6 | 0,278378 | 3,02099 | 0,00252 | 0,097354 | 569,375 | 59,125 | 33,375 |
| pavoniella6 | pavoniella8 | pavoniella4 | 0,278261 | 4,40906 | 1,04E-05 | 0,223464 | 223,375 | 91,875 | 51,875 |
| pavoniella6 | pavoniella7 | pavoniella14 | 0,278182 | 4,49908 | 6,82E-06 | 0,213092 | 195,875 | 87,875 | 49,625 |
| pavonia6 | pavonia9 | pavoniella9 | 0,27771 | 4,05756 | 4,96E-05 | 0,075148 | 173,375 | 131,125 | 74,125 |
| pavoniella8 | pavoniella3 | pavonia3 | 0,277174 | 3,2758 | 0,001054 | 0,074945 | 615 | 58,75 | 33,25 |
| pavoniella13 | pavoniella6 | pavonia5 | 0,27686 | 2,47481 | 0,013331 | 0,057709 | 717,875 | 38,625 | 21,875 |
| pavoniella2 | pavoniella1 | pavonia2 | 0,276817 | 2,55437 | 0,010638 | 0,062305 | 661,625 | 46,125 | 26,125 |
| pavoniella10 | pavoniella5 | pavonia2 | 0,276543 | 3,03676 | 0,002391 | 0,092333 | 559,625 | 64,625 | 36,625 |
| pavoniella8 | pavoniella3 | pavonia10 | 0,276074 | 3,53013 | 0,000415 | 0,063158 | 587,25 | 52 | 29,5 |
| pavoniella9 | pavoniella4 | pavonia2 | 0,276042 | 3,05524 | 0,002249 | 0,08639 | 523,25 | 61,25 | 34,75 |
| pavoniella7 | pavoniella3 | pavonia4 | 0,275964 | 3,30773 | 0,000941 | 0,071926 | 631,75 | 53,75 | 30,5 |
| pavoniella12 | pavoniella11 | pavonia2 | 0,275862 | 2,96475 | 0,003029 | 0,060469 | 609,75 | 46,25 | 26,25 |
| pavoniella13 | pavoniella2 | pavonia5 | 0,275591 | 2,29189 | 0,021912 | 0,058285 | 747,5 | 40,5 | 23 |
| pavonia3 | pavonia7 | pavoniella14 | 0,275532 | 3,79685 | 0,000147 | 0,103187 | 193,625 | 149,875 | 85,125 |
| pavoniella7 | pavoniella11 | pavoniella4 | 0,275441 | 4,44166 | 8,93E-06 | 0,3625 | 150,5 | 117,5 | 66,75 |
| pavoniella3 | pavoniella1 | pavoniella6 | 0,275072 | 4,15156 | 3,30E-05 | 0,206674 | 141,5 | 111,25 | 63,25 |
| pavoniella10 | pavoniella2 | pavoniella4 | 0,27504 | 3,69594 | 0,000219 | 0,318015 | 152,5 | 100,25 | 57 |
| pavoniella13 | pavoniella12 | pavonia9 | 0,274725 | 1,42283 | 0,154785 | 0,0166 | 789,75 | 14,5 | 8,25 |
| pavoniella9 | pavoniella4 | pavonia3 | 0,274038 | 2,98452 | 0,00284 | 0,089063 | 542,75 | 66,25 | 37,75 |
| pavoniella12 | pavoniella11 | pavonia1 | 0,273973 | 2,43086 | 0,015063 | 0,054608 | 625 | 46,5 | 26,5 |
| pavoniella12 | pavoniella10 | pavonia2 | 0,273743 | 2,82755 | 0,004691 | 0,039137 | 644,25 | 28,5 | 16,25 |
| pavoniella13 | pavoniella4 | pavonia1 | 0,273731 | 2,96126 | 0,003064 | 0,085282 | 554,625 | 72,125 | 41,125 |
| pavoniella2 | pavoniella6 | pavonia8 | 0,273684 | 2,98045 | 0,002878 | 0,06762 | 639,5 | 60,5 | 34,5 |
| pavoniella8 | pavoniella2 | pavoniella3 | 0,273342 | 4,15149 | 3,30E-05 | 0,350694 | 141,375 | 117,625 | 67,125 |
| pavoniella9 | pavoniella11 | pavoniella14 | 0,273239 | 3,40155 | 0,00067 | 0,343363 | 132,5 | 113 | 64,5 |
| pavoniella6 | pavoniella9 | pavoniella14 | 0,272727 | 3,68489 | 0,000229 | 0,183499 | 185,75 | 75,25 | 43 |
| pavoniella3 | pavoniella11 | pavoniella6 | 0,272487 | 4,64186 | 3,45E-06 | 0,213472 | 157,75 | 120,25 | 68,75 |
| pavoniella13 | pavoniella1 | pavonia2 | 0,272414 | 2,89912 | 0,003742 | 0,0625 | 604,125 | 46,125 | 26,375 |
| pavoniella10 | pavoniella11 | pavoniella3 | 0,272346 | 4,65153 | 3,29E-06 | 0,35649 | 115,375 | 113,875 | 65,125 |
| pavoniella12 | pavoniella3 | pavonia1 | 0,272237 | 3,07517 | 0,002104 | 0,071733 | 577,25 | 59 | 33,75 |
| pavoniella7 | pavoniella6 | pavonia2 | 0,272085 | 2,0588 | 0,039513 | 0,056995 | 679,75 | 45 | 25,75 |
| pavonia3 | pavonia7 | pavoniella5 | 0,271881 | 4,80077 | 1,58E-06 | 0,105263 | 206,25 | 170,75 | 97,75 |
| pavoniella1 | pavoniella14 | pavonia3 | 0,27182 | 3,75238 | 0,000175 | 0,089271 | 571,75 | 63,75 | 36,5 |
| pavonia8 | pavonia7 | pavoniella5 | 0,271784 | 3,82546 | 0,000131 | 0,092449 | 249,5 | 153,25 | 87,75 |
| pavoniella12 | pavoniella7 | pavoniella6 | 0,271403 | 4,25033 | 2,13E-05 | 0,234646 | 121,75 | 87,25 | 50 |
| pavoniella12 | pavoniella11 | pavoniella5 | 0,270777 | 4,42704 | 9,55E-06 | 0,316119 | 119,5 | 118,5 | 68 |
| pavoniella2 | pavoniella14 | pavonia5 | 0,270627 | 2,42473 | 0,01532 | 0,076279 | 679,375 | 48,125 | 27,625 |
| pavoniella10 | pavoniella4 | pavonia5 | 0,27027 | 3,47179 | 0,000517 | 0,099108 | 615,75 | 58,75 | 33,75 |
| pavoniella11 | pavoniella5 | pavonia3 | 0,269608 | 3,06191 | 0,002199 | 0,083333 | 648,25 | 64,75 | 37,25 |
| pavoniella13 | pavoniella6 | pavonia6 | 0,269231 | 2,05101 | 0,040266 | 0,057377 | 641,125 | 37,125 | 21,375 |
| pavonia5 | pavonia4 | pavoniella9 | 0,26893 | 4,59653 | 4,30E-06 | 0,061438 | 214 | 121,5 | 70 |
| pavoniella6 | pavoniella12 | pavoniella3 | 0,268868 | 4,31052 | 1,63E-05 | 0,2375 | 145,125 | 100,875 | 58,125 |
| pavoniella10 | pavoniella11 | pavonia4 | 0,268382 | 2,50631 | 0,0122 | 0,056765 | 655,625 | 43,125 | 24,875 |
| pavoniella13 | pavoniella11 | pavonia1 | 0,268371 | 2,40948 | 0,015975 | 0,057065 | 631,625 | 49,625 | 28,625 |
| pavoniella8 | pavoniella1 | pavonia2 | 0,267857 | 2,82508 | 0,004727 | 0,058502 | 627,625 | 44,375 | 25,625 |
| pavoniella10 | pavoniella14 | pavonia2 | 0,267857 | 3,82318 | 0,000132 | 0,088757 | 527,125 | 62,125 | 35,875 |
| pavoniella12 | pavoniella3 | pavonia3 | 0,267806 | 3,04094 | 0,002358 | 0,071646 | 587,375 | 55,625 | 32,125 |
| pavoniella10 | pavoniella4 | pavonia3 | 0,267157 | 3,72433 | 0,000196 | 0,085759 | 547,375 | 64,625 | 37,375 |
| pavoniella5 | pavoniella11 | pavoniella9 | 0,266667 | 3,7871 | 0,000152 | 0,20429 | 123,25 | 118,75 | 68,75 |
| pavoniella7 | pavoniella11 | pavoniella5 | 0,26611 | 4,55426 | 5,26E-06 | 0,332836 | 133,125 | 132,625 | 76,875 |
| pavoniella13 | pavoniella9 | pavonia9 | 0,265896 | 1,7747 | 0,075948 | 0,03168 | 624,125 | 27,375 | 15,875 |
| pavonia5 | pavonia4 | pavoniella1 | 0,265823 | 4,76211 | 1,92E-06 | 0,066879 | 196 | 125 | 72,5 |
| pavoniella9 | pavoniella6 | pavonia6 | 0,265823 | 2,00958 | 0,044476 | 0,059716 | 658,25 | 37,5 | 21,75 |
| pavoniella9 | pavoniella1 | pavonia2 | 0,265455 | 2,38764 | 0,016957 | 0,060531 | 584 | 43,5 | 25,25 |
| pavoniella13 | pavoniella5 | pavonia9 | 0,265356 | 2,21729 | 0,026603 | 0,070039 | 577,125 | 64,375 | 37,375 |
| pavoniella5 | pavoniella1 | pavoniella6 | 0,264743 | 2,94319 | 0,003248 | 0,213996 | 143,5 | 126 | 73,25 |
| pavoniella6 | pavoniella9 | pavoniella1 | 0,264117 | 4,08793 | 4,35E-05 | 0,200553 | 125,25 | 86,75 | 50,5 |
| pavoniella9 | pavoniella8 | pavoniella13 | 0,263858 | 2,996 | 0,002736 | 0,154145 | 82,25 | 71,25 | 41,5 |
| pavonia5 | pavonia4 | pavoniella6 | 0,263529 | 5,12506 | 2,97E-07 | 0,066906 | 212,5 | 134,25 | 78,25 |
| pavoniella7 | pavoniella3 | pavonia3 | 0,263158 | 3,71898 | 0,0002 | 0,070896 | 609,5 | 57 | 33,25 |
| pavoniella13 | pavoniella1 | pavonia10 | 0,262745 | 2,39846 | 0,016464 | 0,049192 | 598 | 40,25 | 23,5 |
| pavoniella7 | pavoniella2 | pavoniella4 | 0,26264 | 3,19729 | 0,001387 | 0,305057 | 174,625 | 112,375 | 65,625 |
| pavoniella8 | pavoniella1 | pavonia10 | 0,261538 | 2,69032 | 0,007138 | 0,049455 | 625 | 41 | 24 |
| pavoniella6 | pavoniella7 | pavoniella1 | 0,26087 | 3,98058 | 6,87E-05 | 0,212824 | 131,75 | 94,25 | 55,25 |
| pavoniella12 | pavoniella2 | pavonia5 | 0,260504 | 2,06111 | 0,039292 | 0,05368 | 723,75 | 37,5 | 22 |
| pavoniella7 | pavoniella11 | pavoniella14 | 0,260028 | 3,63892 | 0,000274 | 0,324698 | 145,375 | 113,875 | 66,875 |
| pavoniella9 | pavoniella6 | pavonia1 | 0,259124 | 2,19916 | 0,027867 | 0,050751 | 656,375 | 43,125 | 25,375 |
| pavoniella12 | pavoniella4 | pavonia1 | 0,259009 | 2,777 | 0,005486 | 0,080702 | 547,875 | 69,875 | 41,125 |
| pavoniella2 | pavoniella3 | pavonia2 | 0,258462 | 3,28245 | 0,001029 | 0,063063 | 639,875 | 51,125 | 30,125 |
| pavoniella13 | pavoniella3 | pavonia9 | 0,25816 | 2,42855 | 0,015159 | 0,058903 | 565,75 | 53 | 31,25 |
| pavoniella13 | pavoniella10 | pavonia10 | 0,257862 | 2,31949 | 0,020368 | 0,030461 | 645,75 | 25 | 14,75 |
| pavoniella9 | pavoniella6 | pavonia7 | 0,257703 | 3,22552 | 0,001257 | 0,074194 | 582,625 | 56,125 | 33,125 |
| pavoniella10 | pavoniella4 | pavonia4 | 0,257453 | 3,79746 | 0,000146 | 0,076551 | 572,75 | 58 | 34,25 |
| pavonia6 | pavonia1 | pavoniella14 | 0,256991 | 4,20974 | 2,56E-05 | 0,077355 | 183,25 | 118 | 69,75 |
| pavoniella7 | pavoniella1 | pavoniella4 | 0,256115 | 4,42919 | 9,46E-06 | 0,326007 | 144,625 | 109,125 | 64,625 |
| pavoniella13 | pavoniella3 | pavonia1 | 0,255754 | 2,92434 | 0,003452 | 0,06993 | 586,125 | 61,375 | 36,375 |
| pavonia5 | pavonia4 | pavoniella14 | 0,255501 | 4,76733 | 1,87E-06 | 0,075126 | 186,375 | 128,375 | 76,125 |
| pavonia4 | pavonia8 | pavoniella13 | 0,254023 | 4,05291 | 5,06E-05 | 0,06105 | 222,125 | 136,375 | 81,125 |
| pavoniella9 | pavoniella4 | pavonia4 | 0,253927 | 2,63873 | 0,008322 | 0,080165 | 562,875 | 59,875 | 35,625 |
| pavoniella7 | pavoniella5 | pavonia1 | 0,253731 | 2,67727 | 0,007422 | 0,079705 | 604 | 73,5 | 43,75 |
| pavonia8 | pavonia7 | pavoniella14 | 0,253378 | 3,61747 | 0,000297 | 0,088933 | 236,875 | 139,125 | 82,875 |
| pavoniella11 | pavoniella14 | pavonia1 | 0,253264 | 3,87651 | 0,000106 | 0,072388 | 576,5 | 60 | 35,75 |
| pavoniella9 | pavoniella11 | pavonia6 | 0,253012 | 2,00507 | 0,044955 | 0,059603 | 614 | 39 | 23,25 |
| pavoniella12 | pavoniella4 | pavonia6 | 0,25266 | 3,18406 | 0,001452 | 0,087719 | 552,875 | 58,875 | 35,125 |
| pavoniella9 | pavoniella3 | pavonia4 | 0,252427 | 2,43916 | 0,014722 | 0,06383 | 586,375 | 48,375 | 28,875 |
| pavoniella13 | pavoniella1 | pavoniella14 | 0,252308 | 3,28228 | 0,00103 | 0,292857 | 124,5 | 101,75 | 60,75 |
| pavoniella8 | pavoniella1 | pavonia4 | 0,251799 | 2,22639 | 0,025988 | 0,055162 | 668,75 | 43,5 | 26 |
| pavoniella12 | pavoniella10 | pavonia4 | 0,251799 | 1,97622 | 0,04813 | 0,027111 | 688,5 | 21,75 | 13 |
| pavoniella10 | pavoniella3 | pavonia4 | 0,251613 | 3,20744 | 0,001339 | 0,0626 | 598 | 48,5 | 29 |
| pavonia6 | pavonia10 | pavoniella14 | 0,251596 | 4,69399 | 2,68E-06 | 0,079372 | 169 | 122,5 | 73,25 |
| pavoniella2 | pavoniella5 | pavonia1 | 0,250646 | 2,21268 | 0,02692 | 0,064753 | 682,25 | 60,5 | 36,25 |
| pavoniella9 | pavoniella1 | pavoniella3 | 0,250379 | 3,17118 | 0,001518 | 0,314286 | 106,25 | 103 | 61,75 |
| pavoniella10 | pavoniella1 | pavoniella4 | 0,250364 | 3,823 | 0,000132 | 0,33463 | 131,875 | 107,375 | 64,375 |
| pavoniella10 | pavoniella14 | pavonia6 | 0,25 | 3,67164 | 0,000241 | 0,090278 | 553,875 | 56,875 | 34,125 |
